# Extraordinarily Precise Nematode Sex Ratios: Adaptive Responses to Vanishingly Rare Mating Options

**DOI:** 10.1101/2021.05.25.445688

**Authors:** Justin Van Goor, Edward Allen Herre, Adalberto Gómez, John D. Nason

**Author notes:** Email addresses.

## Abstract

Sex ratio theory predicts both mean sex ratio and variance under a range of population structures. Here, we compare two genera of phoretic nematodes (*Parasitodiplogaster* and *Ficophagus* spp.) associated with twelve fig-pollinating wasp species in Panama. The host wasps exhibit classic Local Mate Competition: only inseminated females disperse from natal figs, and their offspring form mating pools that consist of scores of the adult offspring contributed by one or a few foundress mothers. In contrast, in both nematode genera, only sexually undifferentiated juveniles disperse, and their mating pools routinely consist of eight or fewer adults. Across all mating pool sizes, the sex ratios observed in both nematode genera are consistently female-biased (~0.34 males), which is markedly less female-biased than is often observed in the host wasps (~0.10 males). In further contrast with their hosts, variances in nematode sex ratios are also consistently precise (significantly less than binomial). The constraints associated with predictably small mating pools within highly subdivided populations appear to select for precise sex ratios that contribute both to the reproductive success of individual nematodes, and to the evolutionary persistence of nematode species. We suggest that some form of environmental sex determination underlies these precise sex ratios.

## Introduction

Sex ratio theory provides testable, quantitative predictions of population and individual adaptations expected under a wide range of selective regimes. Following Darwin (1) and Düsing (2), Fisher (3) argued that natural selection within populations of species with separate sexes should favor equal parental investment in the sexes, generally resulting in a population-wide 1:1 (0.50 males) sex ratio (4). However, Darwin knew that many organisms do not exhibit a 1:1 sex ratio; famously leaving “the problem for future generations” (5). Hamilton (6) recognized that Fisher’s argument underlying the 1:1 ratio implicitly assumed large, panmictic populations, and this assumption does not apply to many organisms. Specifically, many species are characterized by subdivided population structures, with male and female adults typically mating in isolated patches. Only mated females disperse from these patches and typically one or a few foundresses contribute offspring to subsequent mating pools (6–8).

Hamilton and others have extended sex ratio theory that was implicitly developed for panmictic populations and formalized expectations under the conditions of Local Mate Competition (LMC). Under LMC, selection is expected to favor female-biased sex ratios, thereby reducing competition among brothers for mates, and increasing the overall productivity of the patches (6–10). Striking correspondence to specific sex ratio predictions are often observed in organisms with life histories characterized by LMC (e.g., in fig pollinating wasps: 6-12). In addition to exhibiting female-biased population sex ratios, individual foundresses in many species appear to facultatively adjust the sex ratio of their offspring in response to cues reflecting the number of other foundresses contributing broods locally to shared mating pools (9–11, 13). Generally, foundresses that contribute relatively fewer offspring tend to produce relatively more male offspring (9–11, 12–15). The validity of this body of evolutionary theory is firmly established by studies in which different experimentally imposed LMC population structures rapidly produce the corresponding predicted sex ratios (16).

In addition to successful predictions concerning population mean and facultative shifts of sex ratios, it is increasingly recognized that the body of LMC theory also makes a series of subtler predictions concerning sex ratio variance. In large, panmictic populations theoretical considerations suggest that there is little or no selection on the sex ratio of individual broods (and therefore no net selection that is expected to affect population wide variance *per se*). In contrast, under many scenarios with subdivided populations, selection for highly precise brood sex ratios that exhibit strikingly less than binomial variance is both expected and, in a several empirical studies, observed (11–12, 14–15, 17).

Whereas theory can predict what sex ratio means or variances are expected under different selective scenarios, sex determining mechanisms can either promote or constrain what sex ratios can actually be achieved in focal species (7–8, 13–14). For example, with sex determining chromosomes (e.g., XX/XY, as in most mammals, or ZW/ZZ, as in most birds and butterflies, or XX/X0 in most nematodes), the odds that any individual offspring develops as a female or male is often roughly approximated as a coin toss. The usual result is close to a 1:1 offspring sex ratio, both within broods of individual mothers and at a population level (4). In addition, variance around the mean sex ratio is usually roughly binomial. In contrast, haplodiploid sex determination in which fertilized eggs usually develop as females and unfertilized eggs usually develop as males provides much greater variety and flexibility in both observed sex ratios and variances (6–15, 17).

One group of haplo-diploid organisms that have been extensively utilized to develop and test predictions from sex ratio theory are the diverse and relatively host-specific wasps (Agaonidae, Chalcidoidae) that pollinate and reproduce within the inflorescences of the roughly 750 species of figs (*Ficus*, Moraceae) (18). These wasps exhibit life histories that effectively define the LMC scenario (Supplement Figure S1, Table S1). Mated, pollen bearing female wasps disperse to a receptive fig inflorescence or “fig” (technically a syconium) that contains scores to hundreds of uniovulate flowers. Typically, one or a few foundress wasps enter the enclosed structure of the receptive fig, pollinate the flowers, and lay eggs in many of them. Oviposition is usually coupled with the wasp depositing a few drops of fluid that cause the oviposited flowers to transform into galls, within which the wasp larvae develop (19). The offspring of successful foundress wasps mature and emerge within the fig to form a local mating pool that usually consist of scores of adult females and males. After mating, the next generation of mated females disperse to a new fig.

The intimate association of the broods of one or a few individual foundress wasps within a developing fig combines with the thousands of such individual figs on individual fig trees to impose highly subdivided population structures on the wasps. Subdivided structure is most extreme when only one foundress wasp contributes offspring to the mating pool within a fig (single foundress brood), in which case there is complete sib-mating. Theory predicts that maternal reproductive success will be increased by producing only enough sons to successfully mate with the daughters (6, 12, 17). This prediction is well supported, as extremely female-biased sex ratios (5-10% male) are regularly observed. Across species, the observed variance in sex ratio in single foundress broods apparently reflects the frequency of single foundress broods (i.e., the intensity of selection on the sex ratios in these broods, 15, Table S1). Within species, mating pool sex ratios in figs that receive more foundress wasps are generally less female biased, suggesting either facultative shifting of sex ratios by foundresses with shared broods or subtler hard-wired behaviors (e.g., “lay a set number of males and then lay females”) that produce similar patterns. In wasp species with higher average foundress numbers, wasp sex ratios associated with any given number of foundresses also exhibit a less extreme female bias (10–11, Table S1).

Reproducing within the highly subdivided populations provided by the fig wasps and their host figs are two genera of nematodes that have independently colonized the fig-wasp mutualism. These nematodes exhibit species-specific associations with the wasps, which they exploit for dispersal and, in some cases, food. *Parasitodiplogaster* (Order Diplogasterida, Family Diplogastridae) are endoparasites of the wasps, whereas *Ficophagus* (Order Aphelenchida, Family Aphelenchoididae), previously referred to as *Schistonchus* in the New World, consume fig tissues (20–28). Despite this dietary difference, both types of nematode are vectored by dispersing female wasps. As with their host wasps, the adults of *Parasitodiplogaster* and *Ficophagus* exhibit separate male and female sexes, with no evidence for self-fertility (e.g., hermaphroditism or parthenogenic reproduction, 22, 26, 28–29).

Unlike their host fig wasps, in which mated females disperse, only sexually undifferentiated juvenile nematodes disperse. Therefore, the host wasps exhibit a strict sense LMC life cycle while the nematodes do not. In fig wasps, there are usually scores of individuals in a mating pool, and the sex of any one individual wasp will have relatively little effect on the overall brood sex ratio. In contrast, nematodes mating pools usually consist of 4 to 10 adults (20, 23–25, 29), and the sex of any individual strikingly affects the overall sex ratio. In fig wasps, the sex ratio expressed in brood sex ratios are primarily determined by a mated foundress’ tendency to fertilize eggs thereby producing daughters (by controlling the exposure of eggs to sperm stored in her spermatheca (9–11, 13–14)). In contrast, each individual nematode develops as either a female or a male only after phoretic dispersal (28). These contrasts between the wasps and the nematodes motivate the following study.

Here, we present field data on mating pool sex ratios from *Parasitodiplogaster* and *Ficophagus* nematode species associated with fig pollinating wasps in twelve Panamanian *Ficus* host species. We find that nematodes exhibit female-biased sex ratios that are consistent and precise (variances are consistently and dramatically lower than binomial). This precision promotes reproductive and transmission success in the face of constraints imposed by consistently small mating pools. We suggest that some form of post-dispersal environmental sex determination (ESD) provides the mechanism underlying these observations.

## Study Taxa and Methods

Between December 2018 and April 2019, we sampled recently pollinated figs of twelve *Ficus* species growing in the proximity of Barro Colorado Island, Republic of Panama. The host fig species that were sampled include: *F. citrifolia*; *F. colubrinae*; *F. costaricana*; *F. dugandii*; *F. nymphaefolia*; *F. obtusifolia*; *F. perforata*; *F. pertusa*; *F. popenoei*; *F. trigonata*; *F.* near *trigonata*; and *F. yoponensis* (30). The corresponding pollinating wasp species belong to the two genera: *Pegoscapus* and *Tetrapus*. They are described as: *Pegoscapus tonduzii*; *P. insularis/orozscoi*; *P. estherae; P*. *longiceps*; *P. piceipes*; *P. hoffmeyerii*; *P. insularis*; *P*. *silvestris*; *P. gemellus*; *P. grandi*; *P. lopesi*; and *Tetrapus ecuadoranus*, respectively (48–50). Previous studies of *Parasitodiplogaster* and *Ficophagus* (COI and 28S rDNA sequences) indicate distinct species associated with each host fig and pollinator wasp species (22, 25–26, 28, 33–34). *Parasitodiplogaster* emerging from *Pegoscapus* wasps have been preserved in amber from the Dominican Republic, confirming a successful evolutionary history for both the nematode and wasp host of at least 22-25 million years (22).

Sexually immature juvenile nematodes disperse from their natal fig with mated adult female wasps. If a wasp carrying *Parasitodiplogaster* or *Ficophagus* nematodes successfully disperses to enter a receptive fig, after the foundress wasp pollinates, oviposits and dies, adult nematodes will emerge from her body. The individual male and female nematodes aggregate to form a mating pool, with eggs laid within the fig (Figure S1). Nematode development is synchronized with wasp development so that as adult wasps begin to emerge from their galls, scores to hundreds of juvenile nematodes that have molted into the infective-stage perform host-seeking nictation behavior by waving back and forth. Only a small fraction of the juvenile nematodes will successfully attach themselves to a female wasp and enter her body (35). Those that do not contact a female pollinating wasp will not survive to reproduce.

The number of infective juvenile nematodes per *individual* foundress pollinator reflects overall nematode density in the natal fig, and successful nematode contacts with a newly emerged wasp. Nematode infection levels per wasp average 4-8 in successfully dispersed foundress wasps (20, 22–25, 27, Figure 1). However, wasps infected with high numbers of nematodes (> 10) are unlikely to successfully disperse to a new fig (25, 27), thereby limiting the size of mating pools in figs (25, 27, 29).

**Figure 1.**
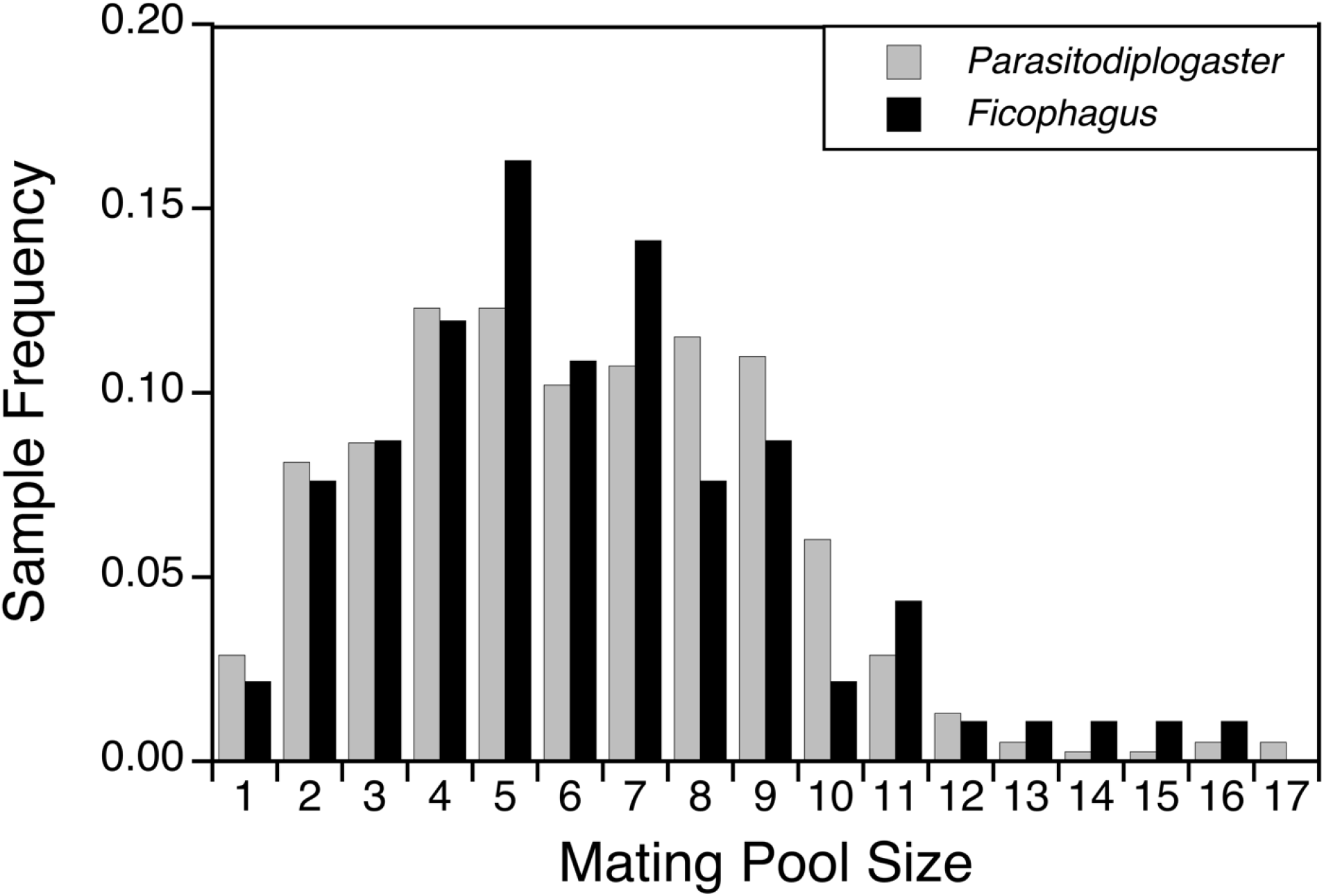
Relative frequencies of mating pool sizes of *Parasitodiplogaster* and *Ficophagus* nematodes found within single-foundress figs are nearly identical (384 and 93 mating pools, respectively). Data were collected and pooled from twelve Panamanian *Ficus* species, with a mean nematode infection load of 6.214 (*Parasitodiplogaster*) and 6.451 (*Ficophagus*) per individual wasp host.

The number of adult nematodes comprising a *mating pool* is determined by the number of infected foundress wasps that enter a given fig and the number of nematodes they each carry. Across the Panamanian species studied here, foundress wasp numbers per fig characteristically range from very low (≥ 95% single foundress) to relatively high (10-20% single foundress) (23, 36). In fig species dominated by only one foundress wasp, the nematodes in a mating pool will typically be introduced by only a single host wasp that acquired its own infection within its natal fig that also likely had only a single infected foundress. For nematode populations to persist in these extreme cases, any fitness effects on the host wasp must be close to neutral, and transmission from infected foundress wasps to their offspring must be essentially perfect (23–25). Further, as is the case with the host wasps, we expect sib-mating will be common and the overall relatedness of nematodes sharing a mating pool will be high (10). In fig species where figs are typically pollinated by multiple foundresses, the constraints on nematode effects on host fitness and requirements for transmission are relatively much less stringent (23–25). Moreover, in these multi-foundress cases, sib-mating and the relatedness among potential mates will be lower, and the number of nematodes per mating pool will often be larger (10, 23–25).

After determining the foundress number, we cut recently pollinated figs into quarters, rinsed them with distilled water, and collected all nematodes. Based on size and morphology, we differentiated adult *Parasitodiplogaster* from the smaller *Ficophagus* (26). Using light microscopy, we then determined the number and sex of adults in mating pools within individual figs. Males of each species were identified based on size, the typical mating position in relation to females (coiled around the female vulval region with genital papillae expressed), and with observations of genitalia (22, 34). In figs with a single foundress, the number of adult males and females (i.e., mating pool sex ratio) associated with individual infected wasps is easy to determine. Importantly, the proportion of successful foundress wasps that carry an infection can also be estimated unambiguously (23–25). With multiple foundresses, it is less clear how many of the wasps were infected and introduced nematodes into shared mating pools. We therefore focused most analyses on single foundress figs, but also analyzed data from multi-foundress figs.

For both *Parasitodiplogaster* and *Ficophagus*, we used a Generalized Linear Model (GLM) with Poisson errors and a log-link function to evaluate the relationship of host fig species and foundress number per fig on mating pool size (*m*) (*m* ~ *host fig species* + *foundress number*). Focusing on single-foundress figs that contained nematodes, we tested the relationship between fig species and mating pool size (*m* ~ *host fig species*). We also conducted two GLMs with a binomial distribution and a logit-link function. The first tested the relationship between sex ratio (*p*_*m*_, proportion of males) of mating pool size *m* and fig species (*p*_*m*_ ~ *host fig species*).The second tested sex ratio as a function of mating pool size (*p*_*m*_ ~ *m*). All GLM analyses were conducted using JMP® Pro 14 (SAS Institute Inc., Cary, NC, 1989-2019).

We used simulations in R (37) to conduct two types of analyses of the precision of observed sex ratios across the nematode mating pools in single foundress figs with different numbers of total adult nematodes (mating pool size). First, we determined whether the observed variance in sex ratio for a given mating pool size was significantly less than the binomial variance expected with sex determining chromosomes (e.g., XX/XY, ZW/ZZ, XX/XO). The expected value of the binomial variance is *p*_*m*_(1-*p*_*m*_)/*m*, where *m* is the mating pool size and *p*_*m*_ is the overall mean sex ratio in *Parasitodiplogaster* or *Ficophagus* (0.315 and 0.353, respectively). We simulated the distribution of the binomial variance in sex ratio for a given mating pool size *m*= 1 to 10, by drawing 10,000 replicates of male-female combinations from the binomial (*m*, *p*_*m*_) with replacement, and calculated the variance in sex ratio across the *n* samples. We conducted a one-tailed test of the hypothesis that the observed variance was less than binomial variance, concluding significance at the 0.05-level if the observed variance was less than or equal to the value of the 500^th^ lowest simulated variance. R code for this simulation is in Supplemental Appendix 1. This simulation provides the expected sex ratio distributions if the sex of individual nematodes is fixed prior to dispersal from their natal fig.

Second, we determined whether the most commonly observed combination of a particular number of males and females (the dominant male-female combination) was observed more frequently than expected given sex determining chromosomes. For mating pool sizes *m* = 1 to 10, we replicated 10,000 samples of observed sample size *n* from the binomial (*m*, *p*_*m*_) with replacement and for each replicate determined the frequency of the dominant male-female combination that was found in the actual data. The one-tailed test of the hypothesis the observed dominant frequency is greater than expected under genetic sex determination was considered significant at the 0.05-level if the observed dominant frequency was less than or equal to the value of the 500^th^ highest simulated dominant frequency. R code for this simulation is in Supplemental Appendix 2. This provides an additional, more explicit test of whether nematode sex is determined prior to the dispersal of juvenile nematodes.

Finally, we further explored the precision of observed sex ratios for both *Parasitodiplogaster* and *Ficophagus* by determining whether the most commonly observed mating pool sex ratios were the closest possible to the observed overall mean sex ratios, given the constraints that small number impose on ratios.

## Results

Of 1668 figs collected from the twelve *Ficus* study species, 915 (55%) showed infections by one or both nematode genera. *Parasitodiplogaster* nematode infections occurred in all twelve sampled *Ficus* species, while *Ficophagus* infections occurred in nine. We determined the number and sex of adult nematodes comprising mating pools for both *Parasitodiplogaster* and *Ficophagus* (523 and 165 figs respectively). The number of individual adult nematodes per mating pool ranged from 1 to 45 and varied significantly with the number of foundresses per fig and the host fig species: *Parasitodiplogaster* (GLM, *n =* 523, *df* = 12, Chi-Square = 597.095, both *p-*values < 0.001) and *Ficophagus* (GLM, *n* = 165, *df =* 9, Chi-Square = 102.026, both *p*-values < 0.001).

Of the 593 figs with detailed information on mating pool composition, 413 were pollinated by a single wasp foundress, indicating that all nematodes involved in each mating pool arrived together with a single wasp. For both genera the median nematode infection per wasp host was 6, with a mean of 6.214 *Parasitodiplogaster* adults (*n* = 384 figs, standard error = 0.158) and a mean of 6.451 *Ficophagus* adults (*n* = 93 figs, standard error = 0.395) per infected wasp (Figure 1). However, for both *Parasitodiplogaster* and *Ficophagus* there was significant heterogeneity in the numbers of nematodes per infected wasp (GLM, *n* = 384, *df* = 11, Chi-Square = 83.630, *p* < 0.001, and GLM, *n* = 93, *df* = 7, Chi-Square = 17.700, *p* = 0.013, respectively). Mating pool sizes are summarized for single-foundress figs in Figure 1, and Table S2. Data from multiple foundress figs are presented in Figure S2 and Table S4.

Across mating pool sizes, nematode sex ratios (*p*_*m*_) in single foundress figs were consistently female-biased for both nematode genera, with a mean sex ratio in *Parasitodiplogaster* of 0.315 (*n* = 384, standard error = 0.006) and in *Ficophagus* of 0.353 (*n* = 93, standard error = 0.010) (Table S2A; for multi-foundress figs see Table S4). GLM analyses indicate that sex ratio, *p*_*m*_, did not vary significantly as a function of host fig species in *Parasitodiplogaster* (GLM, *n* = 384, *df* = 11, Chi-Square = 1.662, *p* = 0.999) or *Ficophagus* (GLM, *n* = 93, *df* = 7, Chi-Square = 0.411, *p* = 0.999). Therefore, in further analyses of sex ratio, single foundress brood sex ratios were pooled across hosts. Similar results were observed in the multi-foundress dataset (both *p*-values > 0.900).

*Parasitodiplogaster* and *Ficophagus* exhibited similar patterns of mating pool composition in single-foundress figs (Tables S2 and S3). In all 13 cases with only one adult nematode, that individual nematode was a female. In 26 of 38 cases of two nematode mating pools there was one male and one female, with 10 cases of two females. In 36 of 41 cases of three nematode mating pools there were two females and one male. In 41of 58 cases of four nematode mating pools there were three females and one male, with 13 cases of two females and two males. Across all mating pool sizes with *m* > 1, there was no shift in sex ratio with increasing mating pool sizes for *Parasitodiplogaster* (GLM, *n* = 372, *df* = 1, Chi-square = 0.025, *p* = 0.875) or *Ficophagus* (GLM, *n* = 91, *df* = 1, Chi-square = 0.011, *p* = 0.917). Further, given that roughly 95% or more of the mating pools encountered in single foundress figs exhibited 11 or fewer adult nematodes, we expect that figs exhibiting 12 or more adult nematodes likely represent cases in which more than one wasp introduced nematodes to the mating pool (Figure 1, Figure S2, Table S2). Nonetheless, we observed no difference in sex ratios in mating pools of 10 or fewer adult nematodes associated with only one infected foundress and those with mating pool sizes of 12 or more (almost certainly associated with two or more infected foundresses, in *Parasitodiplogaster*, *n* = 315, *df* = 1, Chi-square = 0.080, *p* = 0.778; in *Ficophagus, n* = 56, *df* = 1, Chi-square = 0.710, *p* = 0.399).

Importantly, in single foundress figs the observed sample sex ratios across all mating pool sizes were very precise. First, sex ratios exhibited significantly less than the binomial variance expected with sex determining chromosomes (Figure 2, Table S2). This was true for all mating pool sizes in *Parasitodiplogaster*. In *Ficophagus*, where the overall sample sizes were smaller, this pattern was either significant (p < 0.05) or trending (p < 0.1). Non-significant results tended to occur at mating pool sizes with the smallest sample sizes. Second, the dominant male-female combinations were observed significantly more frequently than expected given sex determining chromosomes (Figure 3, Table S3). For *Parasitodiplogaster*, this was the case for all mating pool sizes, either assuming a mean sex ratio of *p*_*m*_ = 0.5 or using observed nematode genus means. In *Ficophagus* this was the case for seven of ten mating pool sizes and marginally significant in two more. Third, the dominant male-female combinations were consistently the closest possible to the nematode genus mean observed sex ratio across all mating pool sizes in both genera (Figure 4, S3, Table S3). Results from nematode mating pools in figs with two or more foundresses do not differ significantly from those for single foundress figs (Tables S2-4).

**Figure 2.**
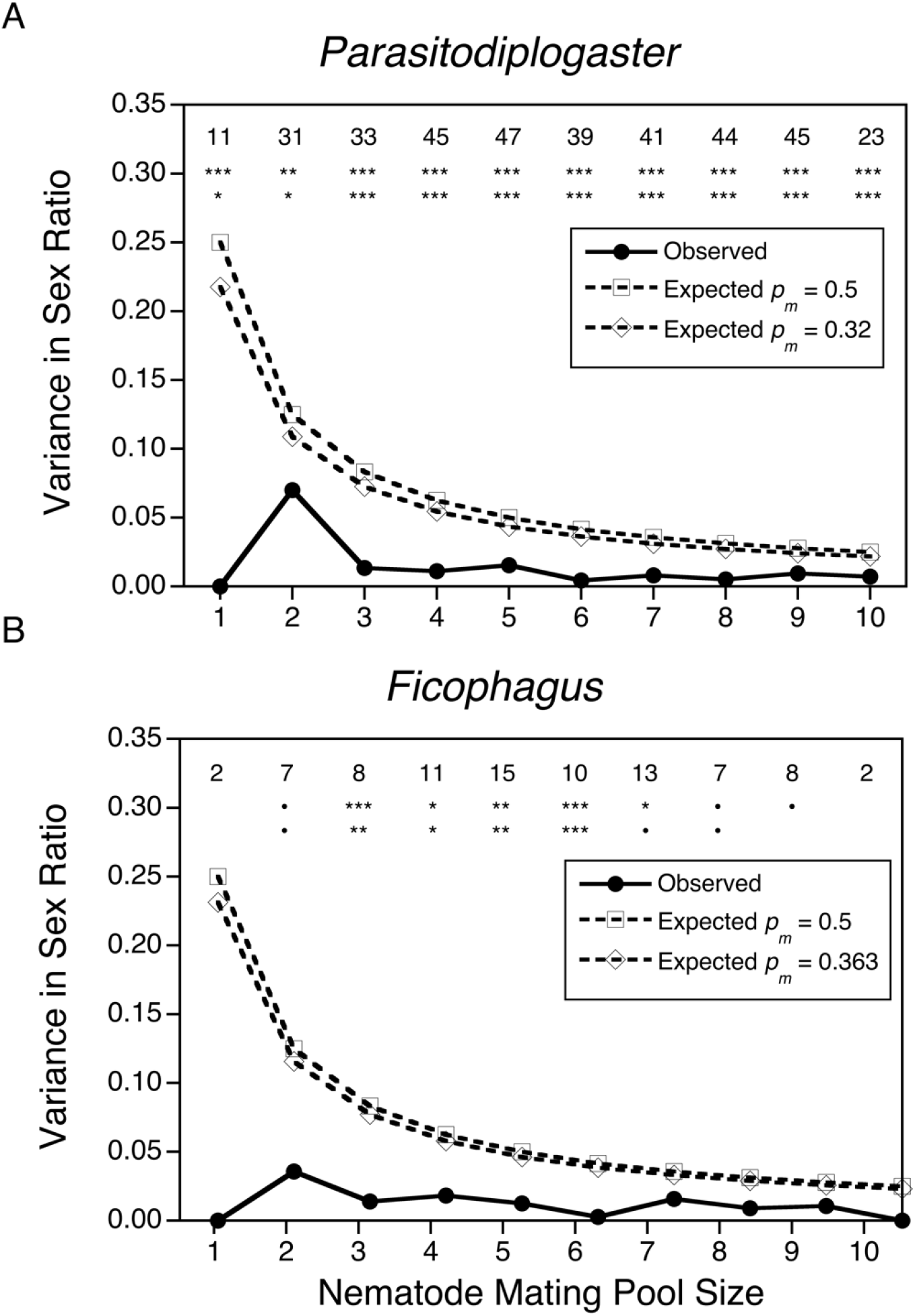
Comparison of the observed (solid line) and expected binomial variance (hatched lines) in sex ratio for *Parasitodiplogaster* (A) and *Ficophagus* (B) mating pools from single foundress figs. The simulated expectations assume binomial sampling given the actual number of mating pools sampled and equal sex ratios (*p*_*m*_ = 0.5) or average sex ratios per genus from single foundress figs (*Parasitodiplogaster*: *p*_*m*_ = 0.315; *Ficophagus*: *p*_*m*_ = 0.353). Observed sample sizes per mating pool size are indicated at the top of panels A and B, followed by significance of the difference between observed and simulated sex ratios assuming *p*_*m*_ = 0.5 and the genus average *p*_*m*_. ***, **, * indicate *p* = < 0.001, *p* = < 0.01 *p* = < 0.05, respectively. • indicates marginal significance (*p* = 0.1 – 0.05). The observed variance is consistently less than binomial expectations.

**Figure 3.**
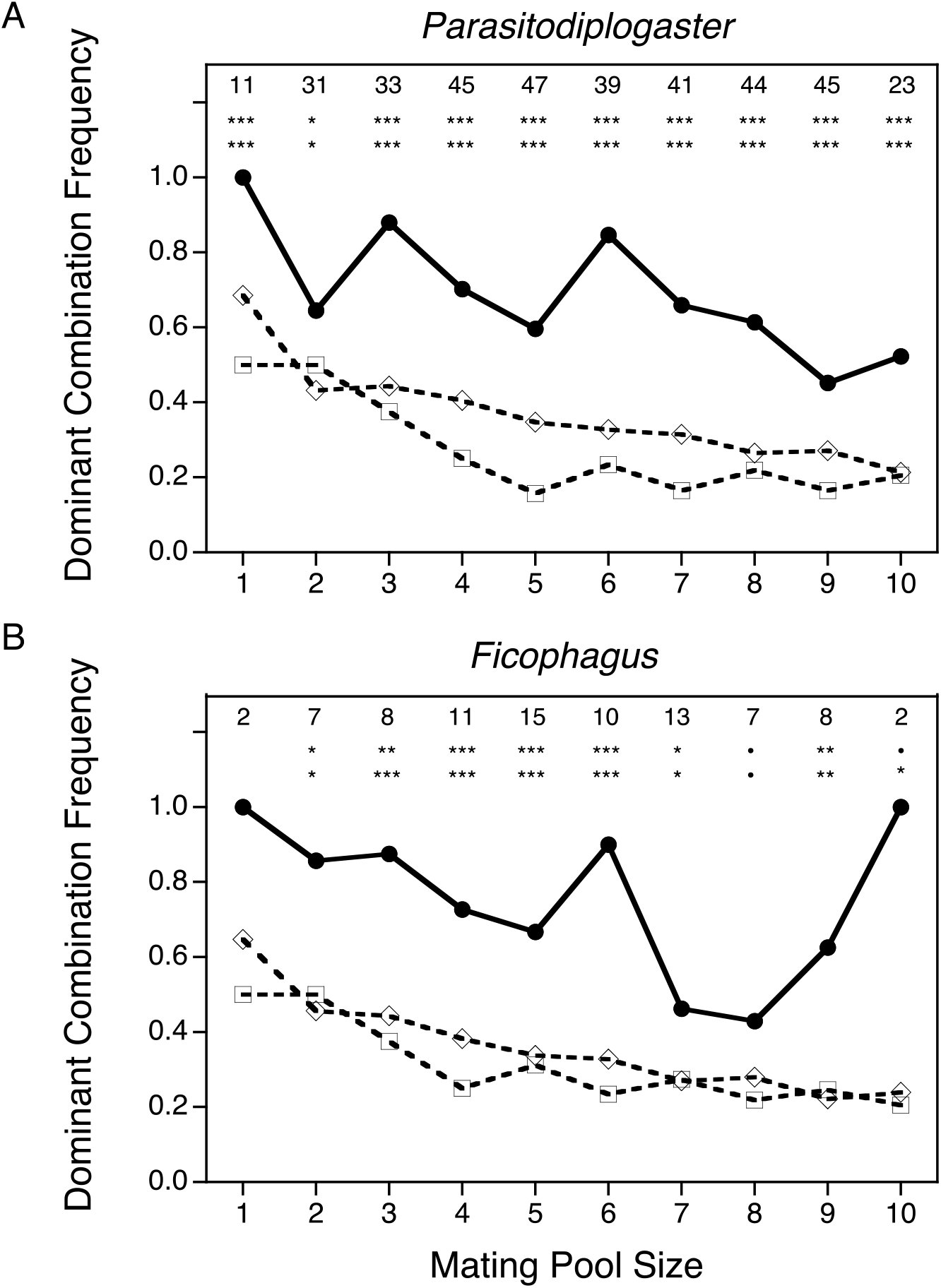
The observed frequency of the dominant (most common) male-female combination (solid lines, with closed circles; see Table 1) for mating pools sampled from single foundress figs (*Parasitodiplogaster* (A) and *Ficophagus* (B)). This frequency is consistently greater than simulated expectations assuming binomial variance given the mating pool size, number of males in the dominant male-female combination, and either equal sex ratios (open squares; *p*_*m*_ = 0.5) or average sex ratios per genus (open diamonds; *Parasitodiplogaster*: *p*_*m*_ = 0.315; *Ficophagus*: *p*_*m*_ = 0.353). Observed sample sizes per mating pool size are indicated at the top of each panel followed by significance of the difference between observed and simulated dominant frequencies for either *p*_*m*_ = 0.5 or the genus average *p*_*m*_. ***, **,* indicate *p* = < 0.001, *p* = < 0.01, *p* = < 0.05, respectively. • indicates marginal significance (*p* = 0.1 – 0.05).

**Figure 4.**
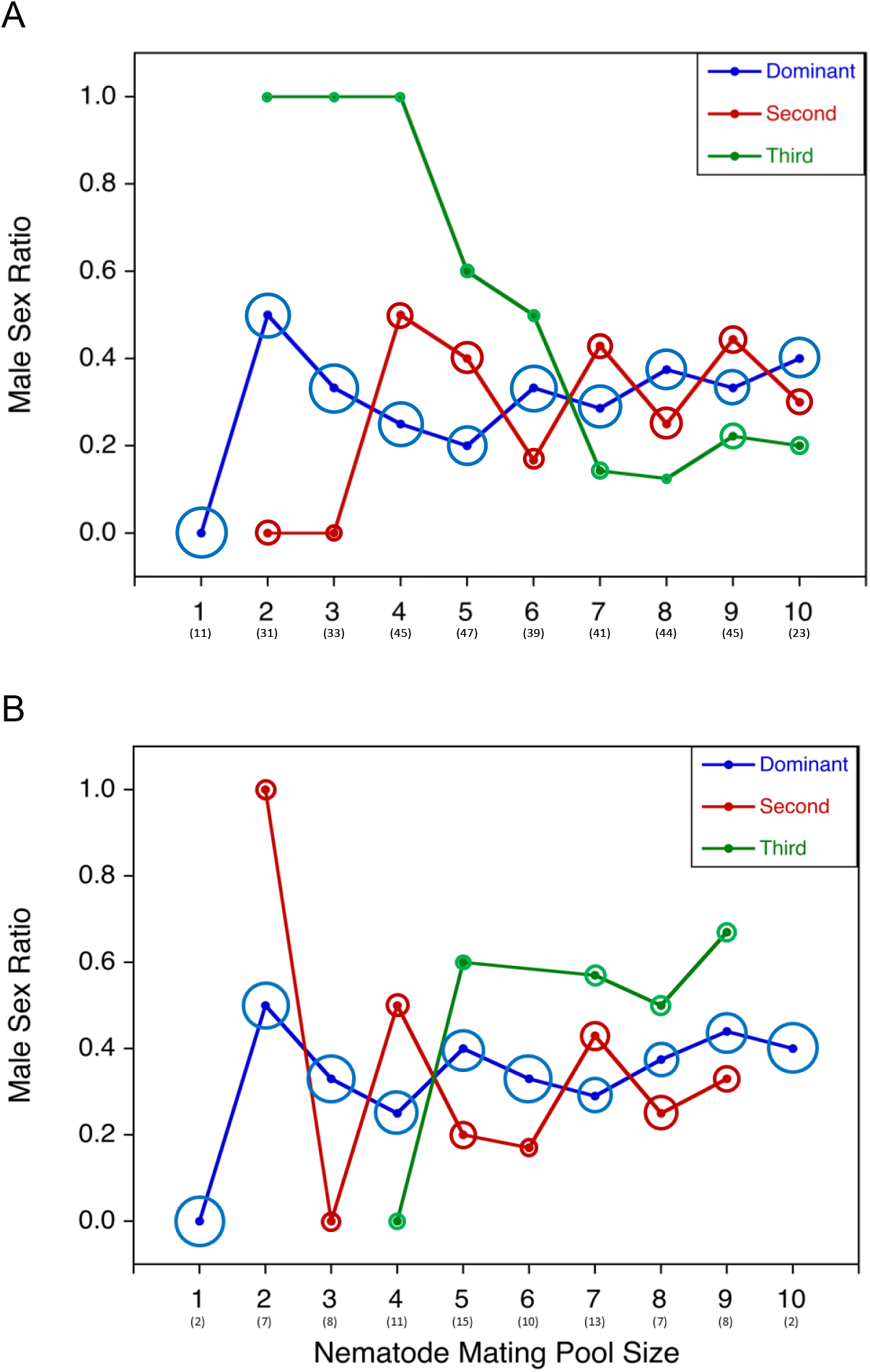
The three most commonly observed male sex ratios (corresponding to specific combinations of males and females) for mating pools ranging from one to ten individuals in single foundress figs (*Parasitodiplogaster* (A) and *Ficophagus* (B)). The sex ratios for the dominant (most common) male-female combination are indicated by the blue datapoints. Given the numerical constraints of small numbers, these are the closest possible to the mean sex ratio for all mating pool sizes greater than one. The second (red) and third (green) most commonly observed sex ratios per mating pool are also presented. The circle surrounding each datapoint correlates to the proportion of data observed for each mating pool size. Sample sizes are given in parentheses beneath each mating pool size.

## Discussion

Two nematode genera (*Parasitodiplogaster* and *Ficophagus*) have independently colonized the fig-pollinator wasp mutualism. In both lineages, there is no evidence of self-fertility (either hermaphroditism or parthenogenesis, 22, 26, 28–29), and only sexually undifferentiated juveniles disperse from their natal fig. Individual nematodes do not mature sexually until after their host fig wasp successfully disperses to a receptive fig where they then form mating pools. Nematode mating pools are regularly composed of a very small number of individual males and females (median 6 total). These aspects of the nematode life cycle contrast with the classical Local Mate Competition (LMC) scenario that characterizes the host wasps (6, 9–11, Table S1). In the wasps, only mated females disperse from mating pools that occur in the natal fig. Further, the mating pools in the hosts consist of scores of adult wasps (36). Across all mating pool sizes, the sex ratios observed in both nematode genera are consistently female-biased (roughly one male for each 2 females, ~0.34 males), which is markedly less female-biased than is often observed in the host wasps (~0.10 males). In further contrast with their hosts (15), variances in nematode sex ratios are also consistently precise (significantly less than binomial), with the most commonly observed sex ratio (combination of males and females) presenting the closest possible sex ratio to the overall mean in both genera. We suggest that constraints associated with predictably small mating pools select for the precise sex ratios, and that some form of Environmental Sex Determination (ESD) provides the mechanism to achieve these precise sex ratios.

Nematode mating pools are regularly composed of a very small number of individual males and females, and, even in the median case of 6 individuals, the sex of any individual nematode has a large effect on the mating pool sex ratio (~0.17). Given predictably small mating pools and apparent lack of self-fertility (22, 26, 28–29), the constraints on nematode sex ratios and sexual reproduction can be most clearly appreciated in cases of the smallest mating pool sizes. If sex in these nematodes were determined as it is in most mammals, birds, or other nematodes, the average sex ratio is expected to be roughly 50% male, with binomial variance. Under these conditions, in mating pools with 2 individual nematodes, ½ of them would consist only of males or females, and represent reproductive dead ends for both nematode individuals. Similarly, in mating pools of 3 or 4 nematodes, ¼ and 1/8 would represent dead ends, respectively. Further, given the observed distributions of mating pool sizes, we would expect 8.5% or more reproductive dead ends, severely threatening the persistence of many of these nematode species (23–25). Instead, across *Parasitodiplogaster* and *Ficophagus,* we found only 2.7% all male or all female (apparent dead end) mating pools (Table S2).

Beyond the scarcity of all female or all male mating pools, the observed nematode sex ratios across all single foundress mating pools sizes are distinctly female biased. All else equal, nematode mating pools with a female bias will be more productive due to more individual females to provide local sources of eggs and juveniles (6–8). Given that higher densities of juvenile nematodes in a fig contributes to the successful infection of departing mated female wasps and, thereby maintain nematode populations in their hosts (23–25), it is not surprising that in *Parasitodiplogaster* and *Ficophagus* the mean observed sex ratios are 0.315 and 0.353 respectively (Tables S2-3). If sex ratio variance were binomial, given the distribution of mating pool sizes we observed, we would expect roughly 33% or more infected figs to exhibit male biases in the nematode mating pools. In contrast, only 1.5% (10 total) of all mating pools exhibited a male bias.

In both nematode genera, the dominant (most common) male-female combinations were also overwhelmingly the closest possible to the overall female biased mean sex ratios, given the ratios possible with the small observed numbers of individuals (~0.34, Figures 3 and 4, Tables S2-4). This pattern in single foundress mating pools was significant in all cases except where mating pool sizes were very small (≤ 2) or sample sizes were unusually low. Specifically, 68% (26 of 38) of two-nematode mating pools exhibited one male and one female nematode (sex ratio = 0.50), with 10 of the remaining 12 two-nematode mating pools exhibiting two female nematodes. This is not consistent with one- and two-nematode mating pools representing random draws from a natal pool with a sex ratio of ~0.34. Similarly, in 88% (36 of 41) of the three-nematode mating pools there were two females and one male. In four-nematode mating pools, the vast majority (97%) of observed mating pools were either one male-three female (41 of 58) or two male-two female (13 of 58) combinations. Similar results were observed for the multi-foundress dataset (Table S4).

Contrasts with the sex ratios exhibited by the Panamanian host wasps are useful for interpreting the potential constraints to the nematodes’ apparently adaptive responses (differences are summarized in Table S1). We have focused our attention on nematode mating pools that resulted from only one wasp foundress introducing adult nematodes into the mating pool. This case is most closely analogous to sex ratios observed in wasp mating pools (broods) from a single foundress. Although in both the wasps and the nematodes the sex ratios are female biased, that bias is much *more* extreme in the wasps (0.05-0.10, in the wasps and ~0.34 in the nematodes, 10–11, 36). Further, the variance in sex ratios is generally much less extreme in the wasps (15, Tables S2-3). We tentatively conclude that the differences between the wasps and the nematodes in both mean and variance of sex ratio can be explained by the extreme differences in their respective mating pool sizes. Moreover, both within and across species, the wasps exhibit less female biased sex ratios in broods with larger numbers of foundresses, (10, 15, 36). In contrast, the nematodes exhibit no evidence for more male biased mating pool sex ratios, either within or across species. The shifts in wasp sex ratios are consistent with adaptive expectations from LMC theory (9–11). Why the nematode mating pools do not analogously exhibit less female biased sex ratios when there is more than one infected wasp contributing adult nematodes, and the mating pool sizes should be sufficient to allow less female biased sex ratios, is not clear. Future experimental and comparative studies that target this question are needed.

Given obligate sexual reproduction and separate sexes, the small number of mating pools (2%) comprised of lone females appear to be reproductive dead ends. This is especially true for nematodes associated with fig species characterized by lower numbers of foundress wasps because fewer adult nematodes are expected per fig. In the fig species with predominantly single-foundress figs, the nematodes must be nearly perfect in transmission from mother to daughter wasps and have negligible negative or even positive effects on wasp fitness (23–25). Persistence on these hosts is especially challenged by the low numbers of adults (4–8) in nematode mating pools, which presumably lower the odds of reproductive success of individual nematodes, as well as threatening both ecological and evolutionary persistence of the species.

At a population level, self-fertile species potentially have up to twice the productivity and a corresponding lower risk of extinction compared to obligately sexual species (6–8, 38–40). Certainly hermaphroditic (41) and parthenogenetic (42–43) nematodes exist. Self-fertility ought to promote higher densities of juvenile nematodes, thereby promoting nematode transmission and local population persistence. Interestingly, the independent evolution of self-fertility has been documented in a few cases within the Diplogastridae (same family as *Parasitodiplogaster*, 44). Yet, neither parthenogenesis nor hermaphroditism has been observed in either *Parasitodiplogaster* or *Ficophagus*.

The extremely stereotyped patterns of numbers of females and males within and across species of both *Parasitodiplogaster* and *Ficophagus* suggest that relatively similar mechanisms influence the sex of maturing nematodes in both groups. Observations in mating pool sizes of one and two are not consistent with a situation in which these nematodes are drawn from an initial pool of two females to each male in the natal fig. Further, the lack of sex ratio shifting in mating pools with more than one infected wasp host suggest that the sex of individuals is most likely determined developmentally within a host wasp just prior to emergence. We suggest that individual nematodes utilize environmental cues (possibly chemical, nutritional, and/or mechanical) that indicate the number and potential sex of other co-occurring nematodes, and influences whether individual juveniles will develop male or female. Here, it appears that this pathway first induces an individual to develop as a female, with subsequent individuals developing as a male. Across mating pool sizes with more than 2 individuals, a relatively constant ratio of females to males (2 to 1) appears to be produced, across species and genera.

A similar ESD-based pattern is found in Mermithid nematodes (Order Mermithida) in which the “founder” or first individual nematode involved in a mating pool usually develops as an adult female, followed by individuals in the pool determining their sex in response to each other (8, 45–47). The phylogenetic divergence between *Parasitodiplogaster*, *Ficophagus*, and Mermithid nematodes suggest that some form of conspecific-influenced ESD mechanism evolved independently and with unique genetic/molecular pathways. Analogous density-dependent ESD scenarios to produce adaptive sex ratios have been described for other nematodes (46, 48–49), malarial *Plasmodium* (50), Ctenophores (51), and in *Daphnia* (52), but are typically associated with life histories characterized by self-fertility. We suggest that this or similar ESD mechanisms allow for precise sex ratio allocation exist in numerous organisms sharing similar life histories, but this prediction awaits future testing.

## Conclusion

*Parasitodiplogaster* and *Ficophagus* represent distinct nematode clades that have independently colonized the fig-wasps mutualism. They exploit different resource bases and have persisted in *Ficus* communities for millions of years, despite obligate sexual reproduction and routinely low numbers of female and male individuals in mating pools. Predictably low mating pool sizes appear to have selected for extremely stereotyped, extraordinarily precise sex ratios. The non-binomial, female-biased sex ratios are consistent with some form of environmental sex determination. However, puzzles remain. Why is there no parthenogenic or hermaphroditic reproduction? Why is there no apparent sex ratio shift (less female biased) observed in mating pools that have more than one source of nematodes within individual figs? What are the mechanisms and evolutionary origins of sex determination in these two genera? More detailed work on the mechanisms underlying these remarkably precise sex ratios and constraints on even greater precision are indicated.

## Acknowledgements

The authors thank D. Adams, P. Dixon, R. Giblin-Davis, J. Greeff, E. Haag, N. Kanzaki, C. Jandér, C. Machado, F. Piatscheck, E.G. Leigh., Jr., S. West, H. Kokko, and an anonymous reviewer for comments and suggestions that improved this manuscript. Funding came from the National Science Foundation (award DEB-1556853 to J. Nason, T. Heath, and E.A. Herre) and from the Smithsonian Tropical Research Institute (to E.A. Herre and J. Van Goor). The authors declare no conflicts of interest.

## Data Accessibility Statement

Data and code used within this manuscript will be publically available on the Dryad and Github repositories and will be included within the supplement.

## Author Contribution Statement

J. Van Goor, E. A. Herre, A. Gómez, and J. Nason all developed the conceptual design of the study, collected data, performed analyses, and contributed to the manuscript. A. Gómez collected the data on *Ficophagus*. J. Nason designed and ran simulations predicting binomial expectations. E.A. Herre and J. Nason provided funding. All authors approve the current draft.

## SUPPLEMENTARY MATERIALS

**Supplemental Table 1.**
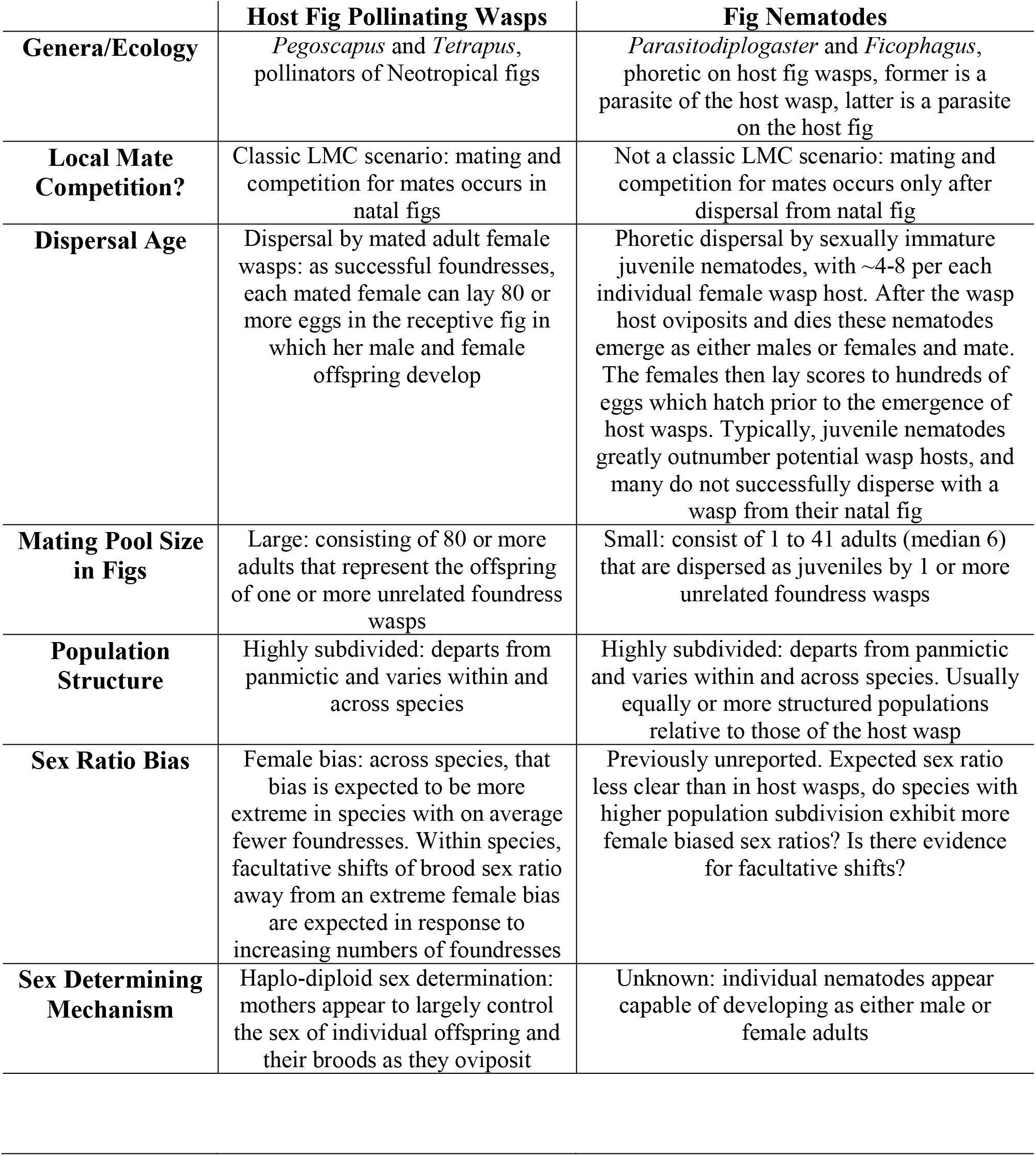
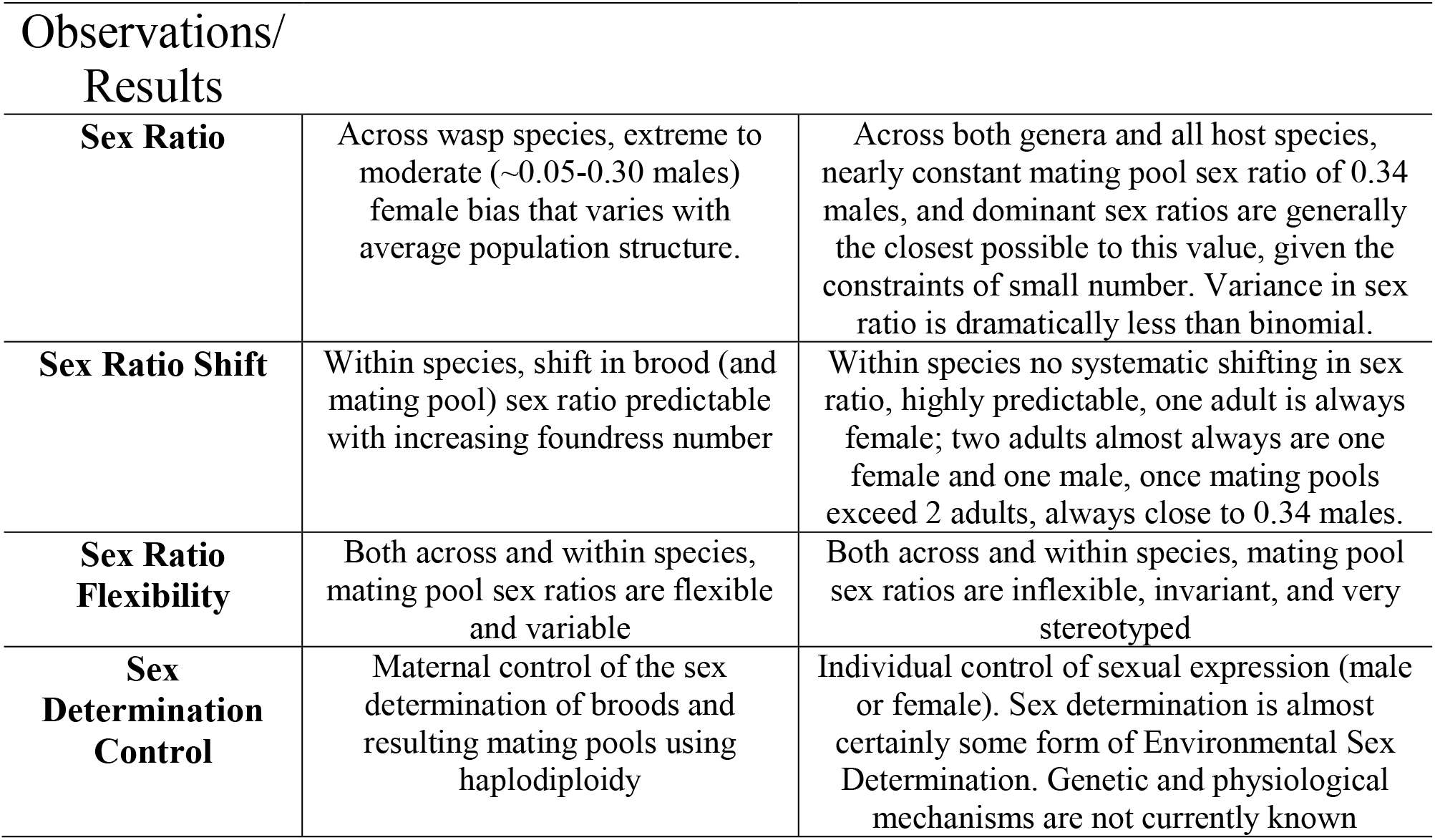
Functional differences between pollinating fig wasp and fig nematode life histories and mating strategies.

|  | **Host Fig Pollinating Wasps** | **Fig Nematodes** |
| --- | --- | --- |
| **Genera/Ecology** | *Pegoscapus* and *Tetrapus*, pollinators of Neotropical figs | *Parasitodiplogaster* and *Ficophagus*, phoretic on host fig wasps, former is a parasite of the host wasp, latter is a parasite on the host fig |
| **Local Mate Competition?** | Classic LMC scenario: mating and competition for mates occurs in natal figs | Not a classic LMC scenario: mating and competition for mates occurs only after dispersal from natal fig |
| **Dispersal Age** | Dispersal by mated adult female wasps: as successful foundresses, each mated female can lay 80 or more eggs in the receptive fig in which her male and female offspring develop | Phoretic dispersal by sexually immature juvenile nematodes, with ~4-8 per each individual female wasp host. After the wasp host oviposits and dies these nematodes emerge as either males or females and mate. The females then lay scores to hundreds of eggs which hatch prior to the emergence of host wasps. Typically, juvenile nematodes greatly outnumber potential wasp hosts, and many do not successfully disperse with a wasp from their natal fig |
| **Mating Pool Size in Figs** | Large: consisting of 80 or more adults that represent the offspring of one or more unrelated foundress wasps | Small: consist of 1 to 41 adults (median 6) that are dispersed as juveniles by 1 or more unrelated foundress wasps |
| **Population Structure** | Highly subdivided: departs from panmictic and varies within and across species | Highly subdivided: departs from panmictic and varies within and across species. Usually equally or more structured populations relative to those of the host wasp |
| **Sex Ratio Bias** | Female bias: across species, that bias is expected to be more extreme in species with on average fewer foundresses. Within species, facultative shifts of brood sex ratio away from an extreme female bias are expected in response to increasing numbers of foundresses | Previously unreported. Expected sex ratio less clear than in host wasps, do species with higher population subdivision exhibit more female biased sex ratios? Is there evidence for facultative shifts? |
| **Sex Determining Mechanism** | Haplo-diploid sex determination: mothers appear to largely control the sex of individual offspring and their broods as they oviposit | Unknown: individual nematodes appear capable of developing as either male or female adults |
| **Sex Ratio** | Across wasp species, extreme to moderate (~0.05-0.30 males) female bias that varies with average population structure. | Across both genera and all host species, nearly constant mating pool sex ratio of 0.34 males, and dominant sex ratios are generally the closest possible to this value, given the constraints of small number. Variance in sex ratio is dramatically less than binomial. |
| **Sex Ratio Shift** | Within species, shift in brood (and mating pool) sex ratio predictable with increasing foundress number | Within species no systematic shifting in sex ratio, highly predictable, one adult is always female; two adults almost always are one female and one male, once mating pools exceed 2 adults, always close to 0.34 males. |
| **Sex Ratio Flexibility** | Both across and within species, mating pool sex ratios are flexible and variable | Both across and within species, mating pool sex ratios are inflexible, invariant, and very stereotyped |
| **Sex Determination Control** | Maternal control of the sex determination of broods and resulting mating pools using haplodiploidy | Individual control of sexual expression (male or female). Sex determination is almost certainly some form of Environmental Sex Determination. Genetic and physiological mechanisms are not currently known |

**Supplemental Table 2.** The observed and expected binomial variance in nematode sex ratio in mating pools from single-foundress figs (all nematodes are introduced by a single infected wasp). Data for *Parasitodiplogaster* presented in Table 1A (384 sample mating pools) and for *Ficophagus* in Table 1B (93 sample mating pools) were collected from twelve Panamanian *Ficus* species. Columns indicate the mating pool size (*m*, number of adult nematodes) found in each fig, the sample size (*n*), the sex ratio (*p*_*m*_, mean proportion males), the observed variance in sex ratio, and then for two sex ratios (*p*_*m*_ = 0.5 and the observed genus mean), the expected binomial variance and the p-value of the difference between the observed and expected sex ratios. *** indicates *p* = < 0.001, ** indicates *p* = < 0.01, * indicates *p* = < 0.05. (+) and (−) indicate the observed variance was greater than or less than a binomial expectation. A. *Parasitodiplogaster*. B. *Ficophagus*.

| Mating pool size<br>($m$) | Sample size<br>($n$) | Sex ratio<br>($p_m$) | Observed variance in<br>sex ratio | Expected variance at<br>$p_m = 0.5$ | $p$-value given<br>$p_m = 0.5$ | Expected variance at<br>$p_m = 0.320$ | $p$-value given<br>$p_m = 0.320$ |
| --- | --- | --- | --- | --- | --- | --- | --- |
| 1 | 11 | 0 | 0 | 0.250 | *** (-) | 0.218 (-) | * (-) |
| 2 | 31 | 0.354 | 0.070 | 0.125 | ** (-) | 0.109 (-) | * (-) |
| 3 | 33 | 0.313 | 0.013 | 0.083 | *** (-) | 0.072 (-) | *** (-) |
| 4 | 47 | 0.328 | 0.021 | 0.063 | *** (-) | 0.054 (-) | *** (-) |
| 5 | 47 | 0.294 | 0.015 | 0.050 | *** (-) | 0.044 (-) | *** (-) |
| 6 | 39 | 0.325 | 0.004 | 0.042 | *** (-) | 0.036 (-) | *** (-) |
| 7 | 41 | 0.310 | 0.008 | 0.036 | *** (-) | 0.031 (-) | *** (-) |
| 8 | 44 | 0.330 | 0.005 | 0.031 | *** (-) | 0.027 (-) | *** (-) |
| 9 | 42 | 0.323 | 0.010 | 0.028 | *** (-) | 0.024 (-) | *** (-) |
| 10 | 23 | 0.357 | 0.007 | 0.025 | *** (-) | 0.022 (-) | *** (-) |
| >10 | 25 | 0.343 |  |  |  |  |  |

| Mating pool size<br>($m$) | Sample size<br>($n$) | Sex ratio<br>($p_m$) | Observed variance in<br>sex ratio | Expected variance at<br>$p_m = 0.5$ | $p$-value given<br>$p_m = 0.5$ | Expected variance at<br>$p_m = 0.353$ | $p$-value given<br>$p_m = 0.353$ |
| --- | --- | --- | --- | --- | --- | --- | --- |
| 1 | 2 | 0 | 0 | 0.250 | 0.505 (-) | 0.231 | 0.540 (-) |
| 2 | 7 | 0.571 | 0.036 | 0.125 | 0.063 (-) | 0.116 | 0.054 (-) |
| 3 | 8 | 0.291 | 0.014 | 0.083 | *** (-) | 0.077 | ** (-) |
| 4 | 11 | 0.273 | 0.018 | 0.063 | * (-) | 0.058 | * (-) |
| 5 | 15 | 0.360 | 0.013 | 0.050 | ** (-) | 0.046 | ** (-) |
| 6 | 10 | 0.317 | 0.003 | 0.042 | *** (-) | 0.039 | *** (-) |
| 7 | 13 | 0.363 | 0.016 | 0.036 | * (-) | 0.033 | 0.062 (-) |
| 8 | 7 | 0.340 | 0.009 | 0.031 | 0.060 (-) | 0.029 | 0.071 (-) |
| 9 | 8 | 0.444 | 0.011 | 0.028 | 0.096 (-) | 0.026 | 0.122 (-) |
| 10 | 2 | 0.400 | 0 | 0.025 | 0.177 (-) | 0.023 | 0.180 (-) |
| >10 | 10 | 0.349 |  |  |  |  |  |

**Supplementary Table 3.** Summary of the most common (dominant) male-female combinations for nematode mating pools collected from single foundress figs in twelve Panamanian *Ficus* species. Data for *Parasitodiplogaster* are presented in Table 2A (384 sample mating pools, mean sex ratio *p*_*m*_ = 0.315), and for *Ficophagus* in Table 2B (93 sample mating pools, mean sex ratio *p*_*m*_ = 0.353). Columns show the mating pool size (*m*, number of adult nematodes). For each mating pool size, the sample number of figs (*n*), the sex ratio (*p*_*m*_, mean proportion males), the most common (dominant) male-female combination, the frequency of this combination, and the probabilities of this frequency under chromosomal sex determination. These probabilities were simulated for two assumed sex ratios (*p*_*m*_ = 0.5 and the observed genus mean). The observed combination frequencies were always greater (+) than expected if nematodes were drawn from a source pool (e.g., natal fig) in which sex (male or female) chromosomally determined. (***, **, * indicate *p* = < 0.001, = < 0.01, and *p* = < 0.05, respectively. These results are not consistent with nematode sex ratio being determined at the time of juvenile nematode dispersal. A. *Parasitodiplogaster*. B. *Ficophagus*.

| Mating pool size ($m$) | Sample size ($n$) | Sex ratio ($p_m$) | Dominant male-female combination | Dominant combination frequency | Expected frequency at $p_m = 0.5$ | $p$-value given $p_m = 0.5$ | Expected frequency at $p_m = 0.315$ | $p$-value given $p_m = 0.315$ |
| --- | --- | --- | --- | --- | --- | --- | --- | --- |
| 1 | 11 | 0 | 0♂ 1♀ (11 of 11) | 1 | 0.500 | *** (+) | 0.685 | *** (+) |
| 2 | 31 | 0.354 | 1♂ 1♀ (20 of 31) | 0.645 | 0.500 | * (+) | 0.432 | * (+) |
| 3 | 33 | 0.313 | 1♂ 2♀ (29 of 33) | 0.879 | 0.375 | *** (+) | 0.443 | *** (+) |
| 4 | 47 | 0.328 | 1♂ 3♀ (33 of 47) | 0.702 | 0.250 | *** (+) | 0.405 | *** (+) |
| 5 | 47 | 0.294 | 1♂ 4♀ (28 of 47) | 0.596 | 0.156 | *** (+) | 0.347 | *** (+) |
| 6 | 39 | 0.325 | 2♂ 4♀ (33 of 39) | 0.846 | 0.234 | *** (+) | 0.328 | *** (+) |
| 7 | 41 | 0.310 | 2♂ 5♀ (27 of 41) | 0.659 | 0.164 | *** (+) | 0.314 | *** (+) |
| 8 | 44 | 0.330 | 3♂ 5♀ (27 of 44) | 0.614 | 0.219 | *** (+) | 0.264 | *** (+) |
| 9 | 42 | 0.323 | 3♂ 6♀ (19 of 42) | 0.452 | 0.164 | *** (+) | 0.271 | *** (+) |
| 10 | 23 | 0.357 | 4♂ 6♀ (12 of 23) | 0.522 | 0.205 | *** (+) | 0.214 | *** (+) |
| >10 | 25 | 0.343 |  |  |  |  |  |  |

| Mating pool size ($m$) | Sample size ($n$) | Sex ratio ($p_m$) | Dominant male-female combination | Dominant combination frequency | Expected frequency at $p_m = 0.5$ | $p$-value given $p_m = 0.5$ | Expected frequency at $p_m = 0.353$ | $p$-value given $p_m = 0.353$ |
| --- | --- | --- | --- | --- | --- | --- | --- | --- |
| 1 | 2 | 0 | 0♂ 1♀ (2 of 2) | 1 | 0.500 | 0.318 (+) | 0.647 | 0.324 (+) |
| 2 | 7 | 0.571 | 1♂ 1♀ (6 of 7) | 0.857 | 0.500 | * (+) | 0.457 | * (+) |
| 3 | 8 | 0.291 | 1♂ 2♀ (7 of 8) | 0.875 | 0.375 | ** (+) | 0.443 | *** (+) |
| 4 | 11 | 0.273 | 1♂ 3♀ (8 of 11) | 0.727 | 0.250 | *** (+) | 0.382 | ** (+) |
| 5 | 15 | 0.360 | 2♂ 3♀ (10 of 15) | 0.667 | 0.313 | *** (+) | 0.337 | *** (+) |
| 6 | 10 | 0.317 | 2♂ 4♀ (9 of 10) | 0.900 | 0.234 | *** (+) | 0.328 | *** (+) |
| 7 | 13 | 0.363 | 3♂ 4♀ (6 of 13) | 0.462 | 0.273 | * (+) | 0.270 | * (+) |
| 8 | 7 | 0.340 | 3♂ 5♀ (3 of 7) | 0.429 | 0.219 | 0.068 (+) | 0.279 | 0.070 (+) |
| 9 | 8 | 0.444 | 4♂ 5♀ (5 of 8) | 0.625 | 0.246 | ** (+) | 0.222 | ** (+) |
| 10 | 2 | 0.400 | 4♂ 6♀ (2 of 2) | 1 | 0.205 | 0.051 (+) | 0.239 | * (+) |
| >10 | 10 | 0.349 | N/A |  |  |  |  |  |

**Supplemental Table 4A.**
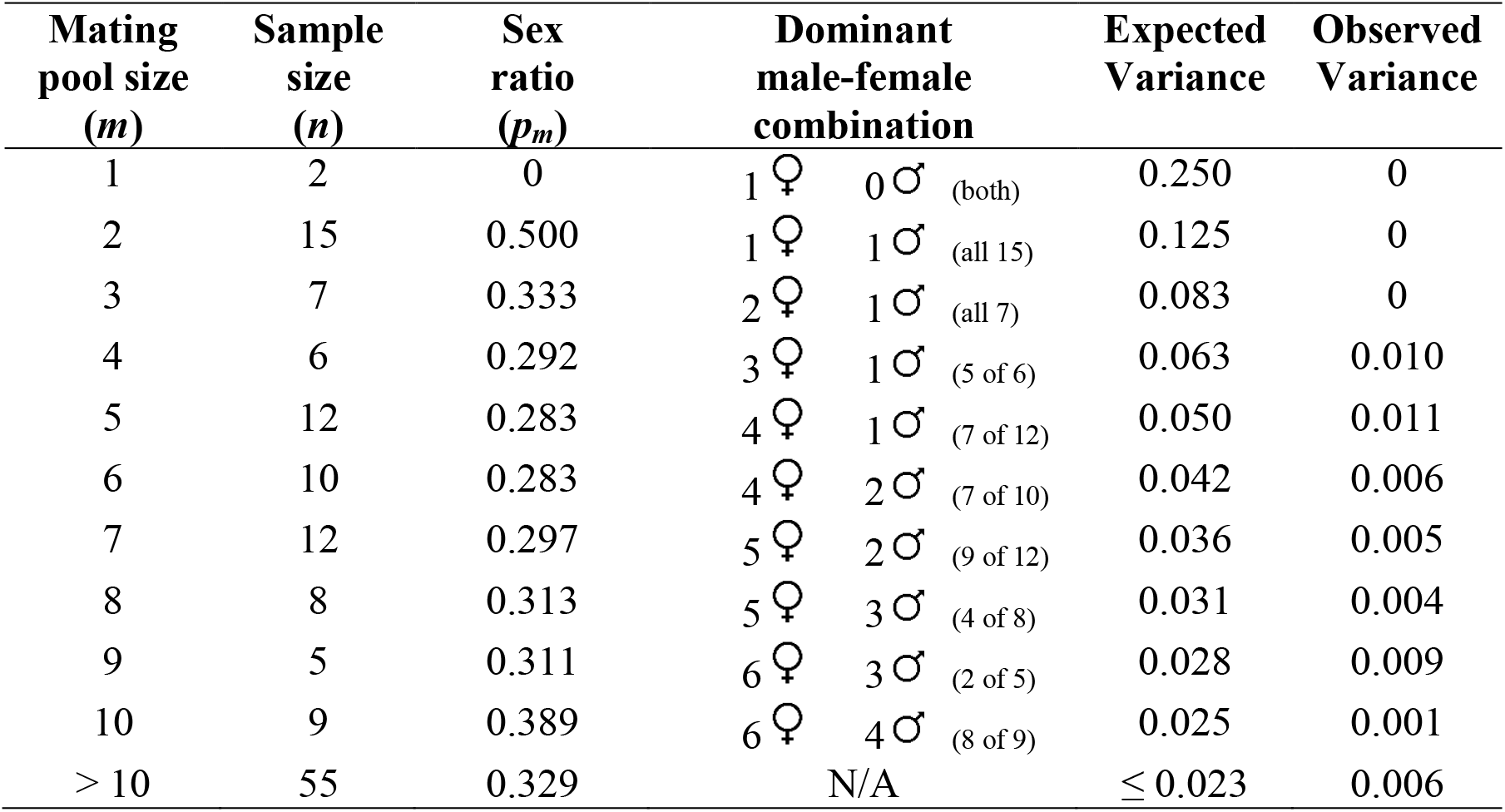
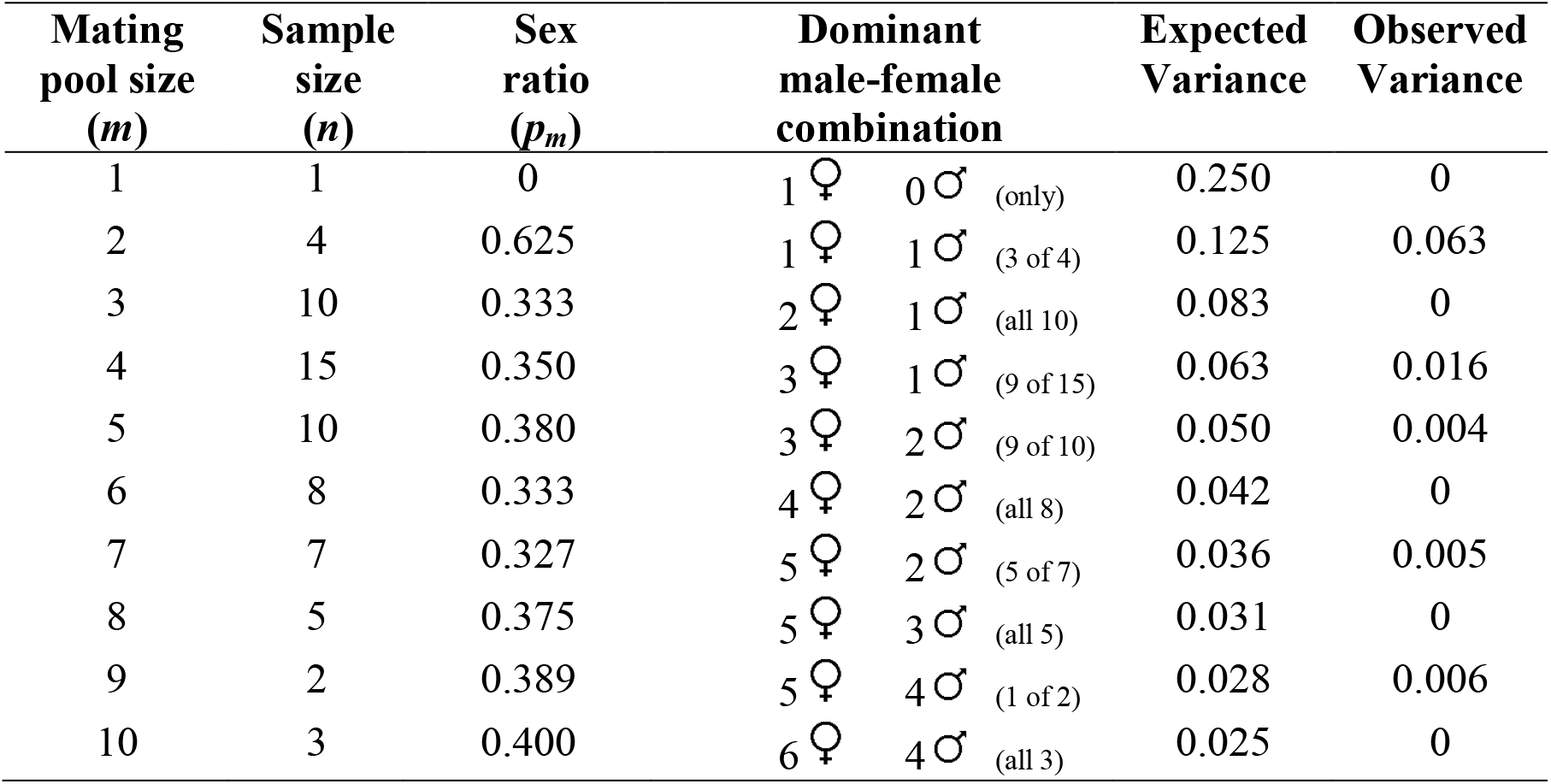
Summary data for nematode mating pools for figs containing two or more foundress. These figs were collected from twelve Panamanian *Ficus* species, with data for *Parasitodiplogaster* presented in Table 1A (141 sample mating pools, mean sex ratio *p*_*m*_ = 0.334) and for *Ficophagus* in Table 1B (85 sample mating pools, mean sex ratio *p*_*m*_ = 0.371). Columns show the mating pool size (*m*, number of adult nematodes) found in each fig, the sample size (*n*), the sex ratio (*p*_*m*_, mean proportion males), the dominant (most common) male-female combination, and the expected and observed variance in sex ratio (expected based on the binomial distribution with *p*_*m*_ = 0.5).

| Mating pool size ($m$) | Sample size ($n$) | Sex ratio ($p_m$) | Dominant male-female combination | Expected Variance | Observed Variance |
| --- | --- | --- | --- | --- | --- |
| 1 | 2 | 0 | 1 ♀ 0 ♂ (both) | 0.250 | 0 |
| 2 | 15 | 0.500 | 1 ♀ 1 ♂ (all 15) | 0.125 | 0 |
| 3 | 7 | 0.333 | 2 ♀ 1 ♂ (all 7) | 0.083 | 0 |
| 4 | 6 | 0.292 | 3 ♀ 1 ♂ (5 of 6) | 0.063 | 0.010 |
| 5 | 12 | 0.283 | 4 ♀ 1 ♂ (7 of 12) | 0.050 | 0.011 |
| 6 | 10 | 0.283 | 4 ♀ 2 ♂ (7 of 10) | 0.042 | 0.006 |
| 7 | 12 | 0.297 | 5 ♀ 2 ♂ (9 of 12) | 0.036 | 0.005 |
| 8 | 8 | 0.313 | 5 ♀ 3 ♂ (4 of 8) | 0.031 | 0.004 |
| 9 | 5 | 0.311 | 6 ♀ 3 ♂ (2 of 5) | 0.028 | 0.009 |
| 10 | 9 | 0.389 | 6 ♀ 4 ♂ (8 of 9) | 0.025 | 0.001 |
| > 10 | 55 | 0.329 | N/A | $\leq 0.023$ | 0.006 |
| 1 | 1 | 0 | 1 ♀ 0 ♂ (only) | 0.250 | 0 |
| 2 | 4 | 0.625 | 1 ♀ 1 ♂ (3 of 4) | 0.125 | 0.063 |
| 3 | 10 | 0.333 | 2 ♀ 1 ♂ (all 10) | 0.083 | 0 |
| 4 | 15 | 0.350 | 3 ♀ 1 ♂ (9 of 15) | 0.063 | 0.016 |
| 5 | 10 | 0.380 | 3 ♀ 2 ♂ (9 of 10) | 0.050 | 0.004 |
| 6 | 8 | 0.333 | 4 ♀ 2 ♂ (all 8) | 0.042 | 0 |
| 7 | 7 | 0.327 | 5 ♀ 2 ♂ (5 of 7) | 0.036 | 0.005 |
| 8 | 5 | 0.375 | 5 ♀ 3 ♂ (all 5) | 0.031 | 0 |
| 9 | 2 | 0.389 | 5 ♀ 4 ♂ (1 of 2) | 0.028 | 0.006 |
| 10 | 3 | 0.400 | 6 ♀ 4 ♂ (all 3) | 0.025 | 0 |

**Supplemental Figure 1.**
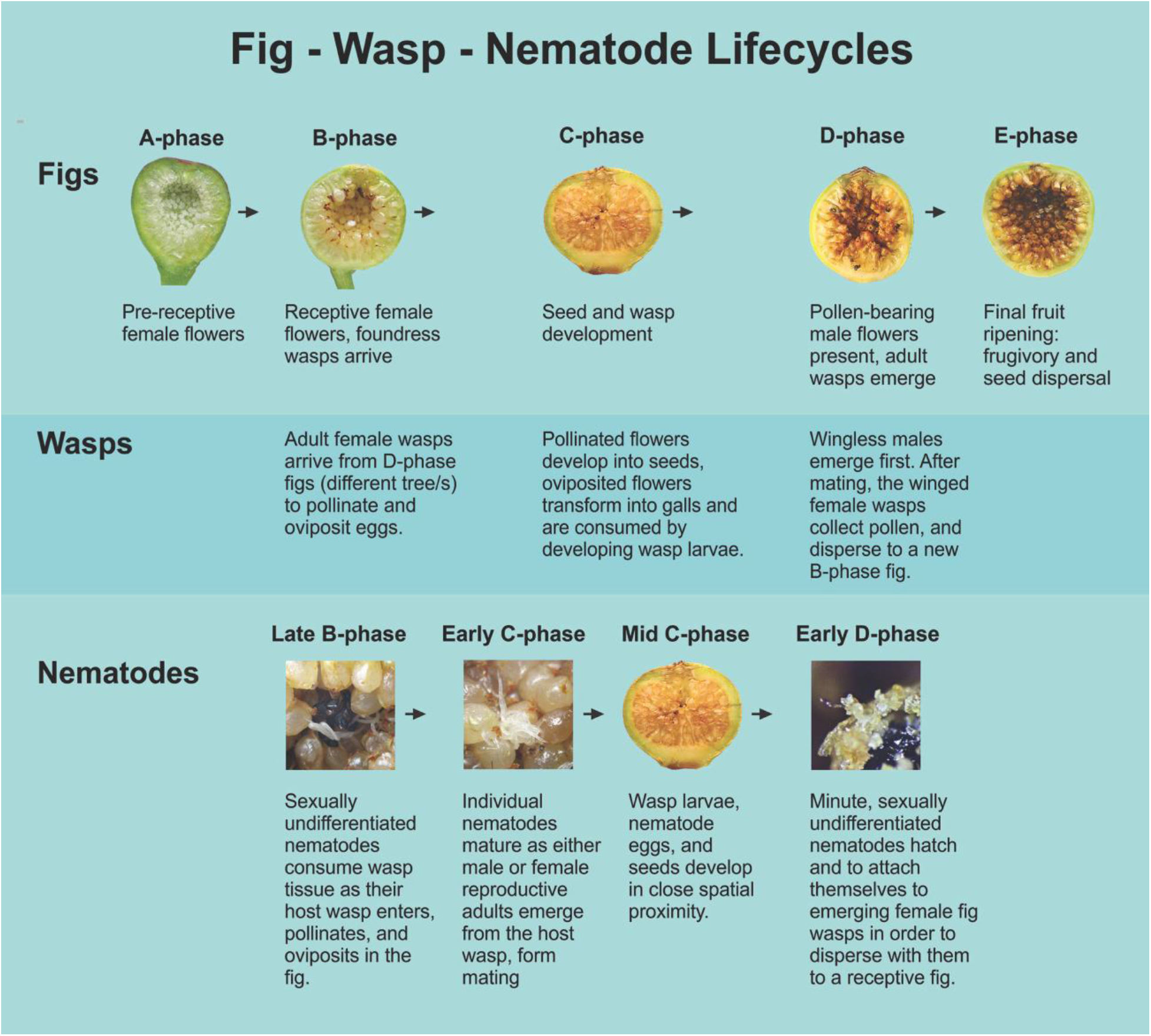
Differences in life history characteristics between pollinating figs wasps and their associated fig nematodes (*Parasitodiplogaster* and *Ficophagus*) within various stages of fig development.

**Supplemental Figure 2.**
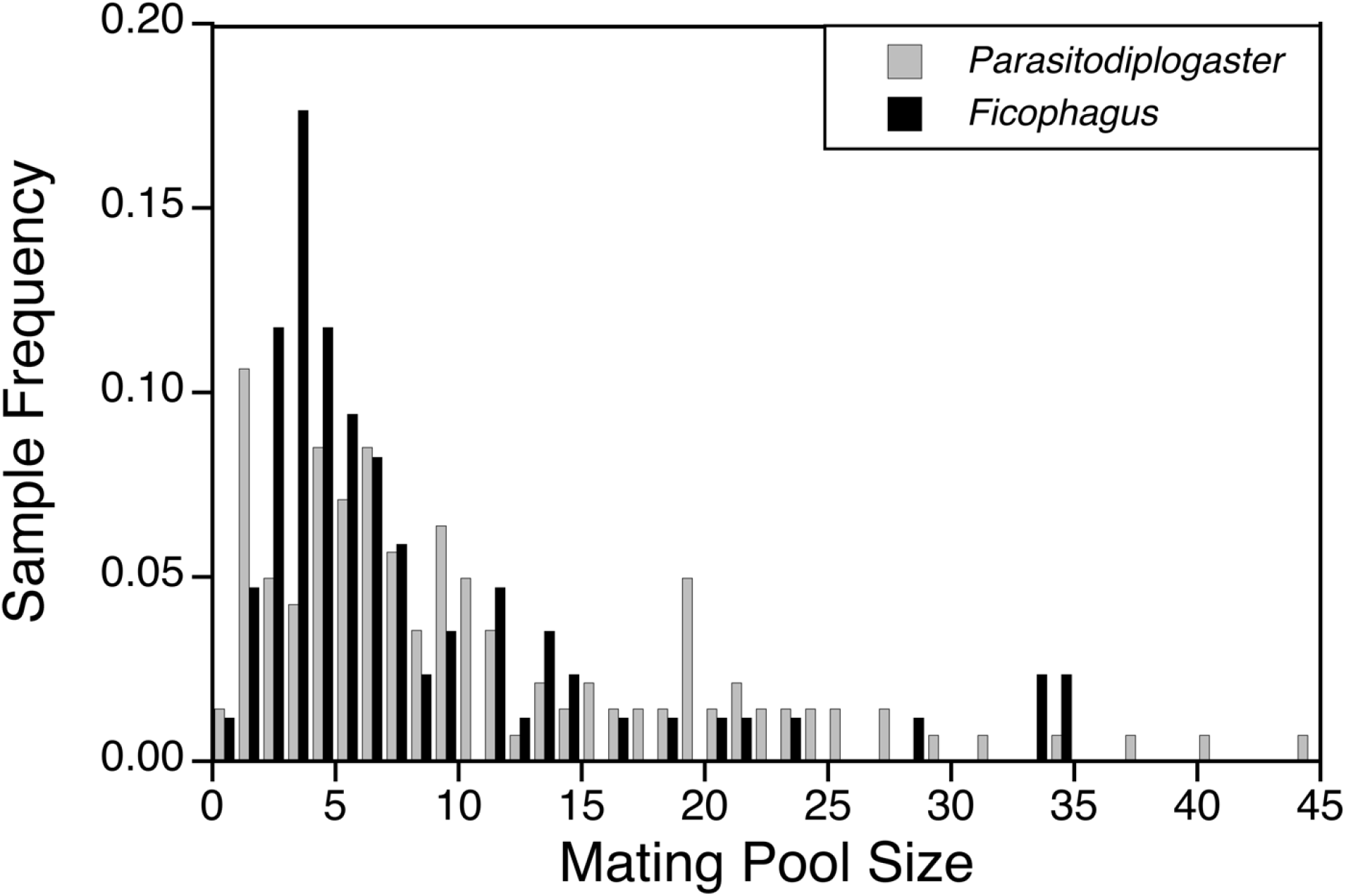
Relative frequencies of *Parasitodiplogaster* and *Ficophagus* sample mating pool sizes (141 and 85 mating pools, respectively) from twelve Panamanian *Ficus* species exhibited in figs with two or more foundresses. The number of nematode individuals observed in these multi-foundress mating pools was greater in *Parasitodiplogaster* (mean = 11.199, median = 8, standard error = 0.743) and *Ficophagus* (mean = 8.671, median = 6, standard error = 0.850) than in single-foundress mating pools (means of 6.214 and 6.451, respectively; Figure 1), suggesting that in at least some cases, more than one foundress wasps introduced a nematode infection to the shared fig.

**Supplementary Figure 3.**
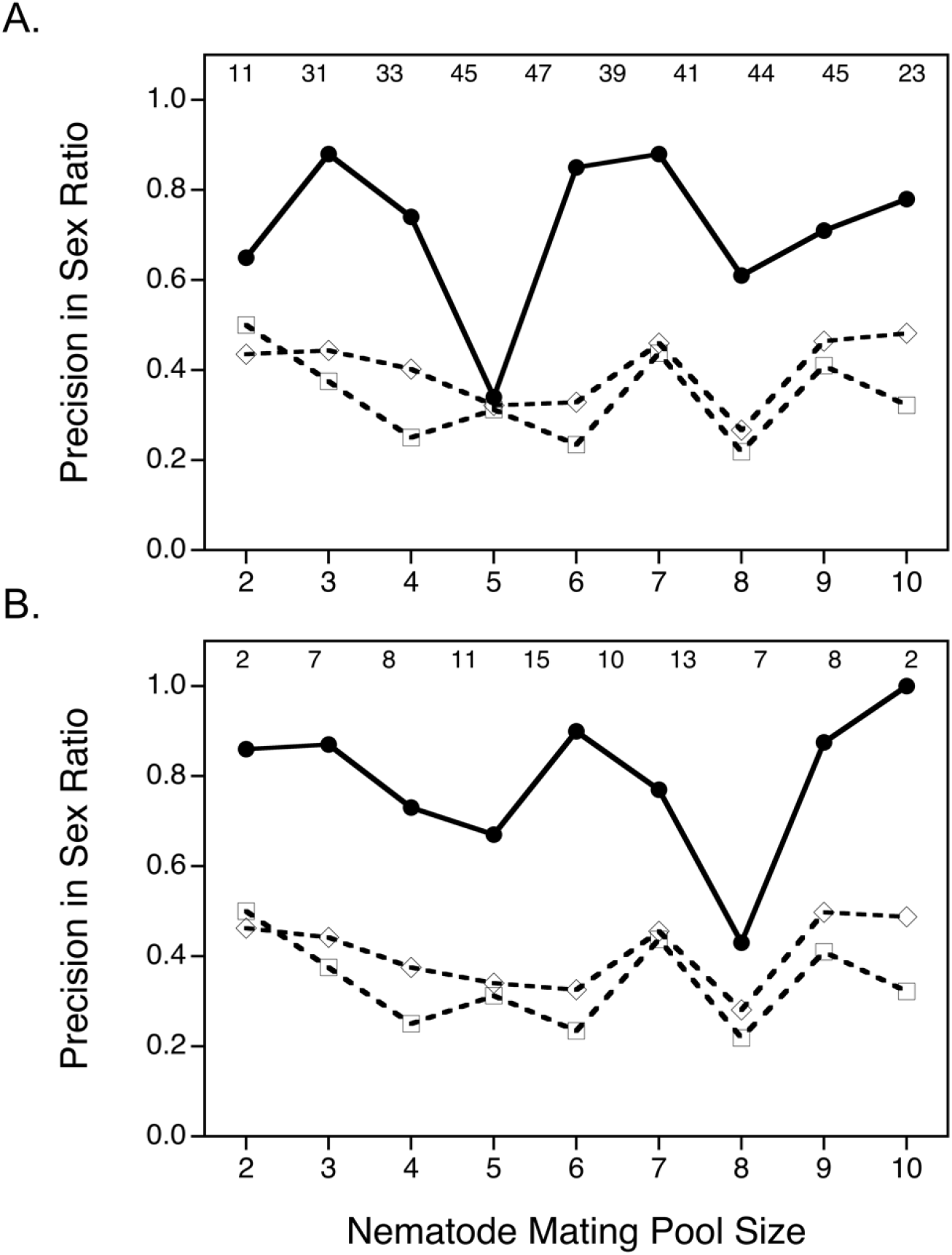
The frequency with which the observed or simulated data exhibit the sex ratio closest to the overall mean (precision) in sex ratio (solid line) in single foundress mating pools for both *Parasitodiplogaster* (A) and *Ficophagus* (B). These are compared to their expected binomial frequencies (expected under chromosomal sex determination) given the actual number of mating pools sampled for both equal sex ratios (*p*_*m*_ = 0.5; dashed line with open squares) and the observed average sex ratios (*Parasitodiplogaster*: *p*_*m*_ = 0.315; *Ficophagus*: *p*_*m*_ = 0.353; dashed line with open diamonds). Observed sex ratios (combinations of male and females) are consistently the closest possible value to the overall mean sex ratio for each genus of nematode, given the constraints imposed by small numbers (mating pool sizes). Observed sample sizes per mating pool size are indicated at the top of each panel.

## Supplemental Appendix 1.

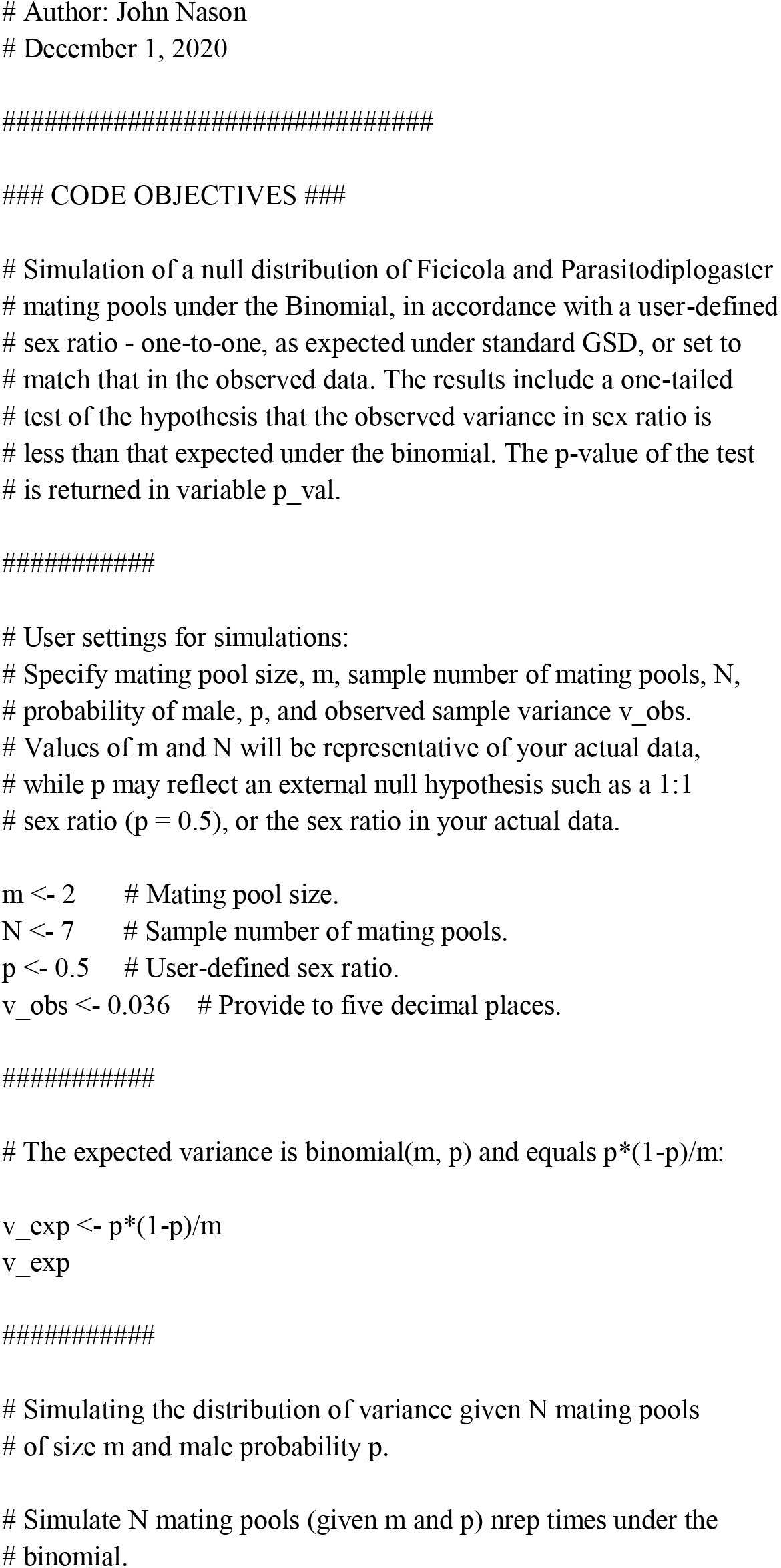
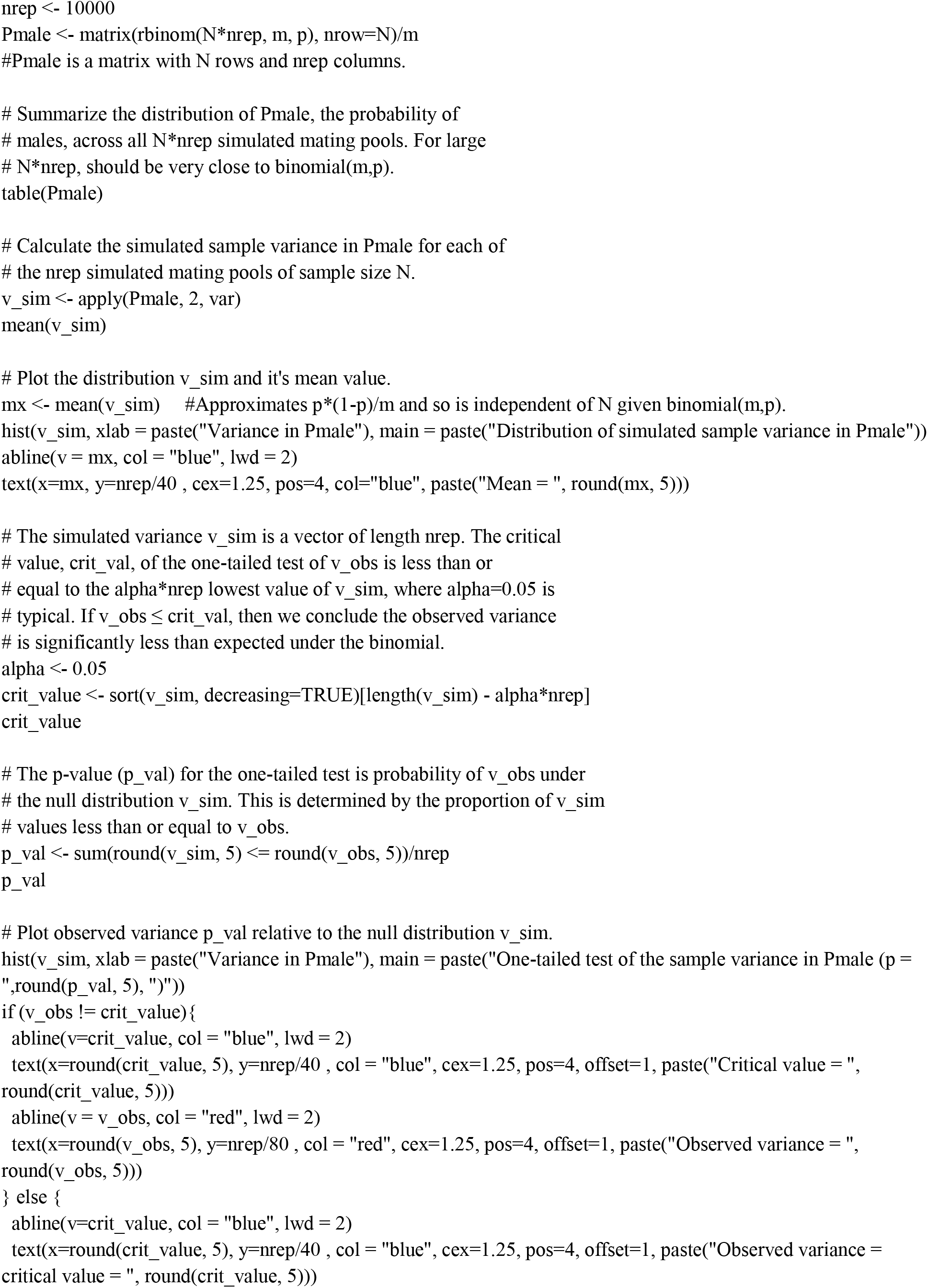
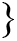

## Supplemental Appendix 2.

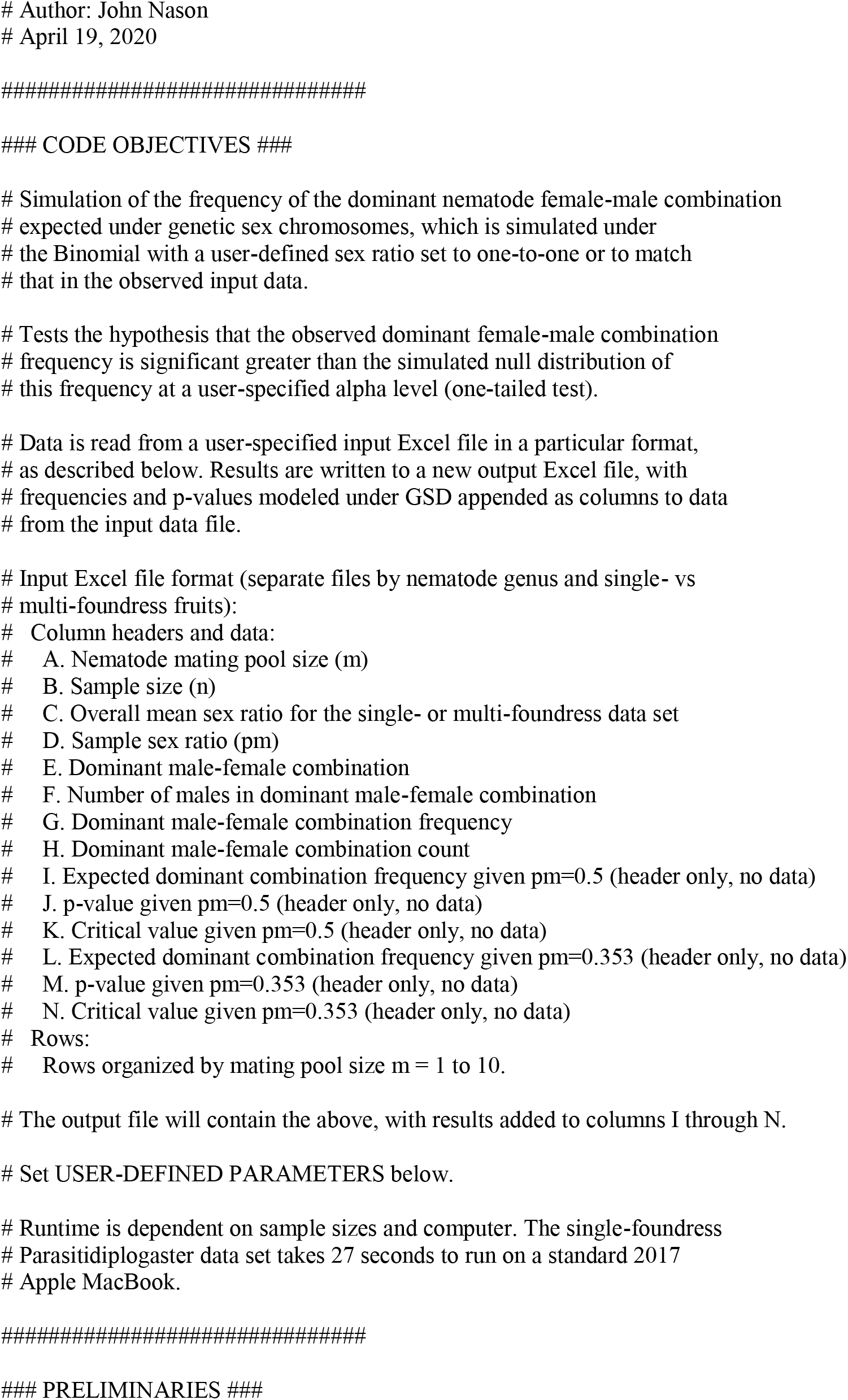
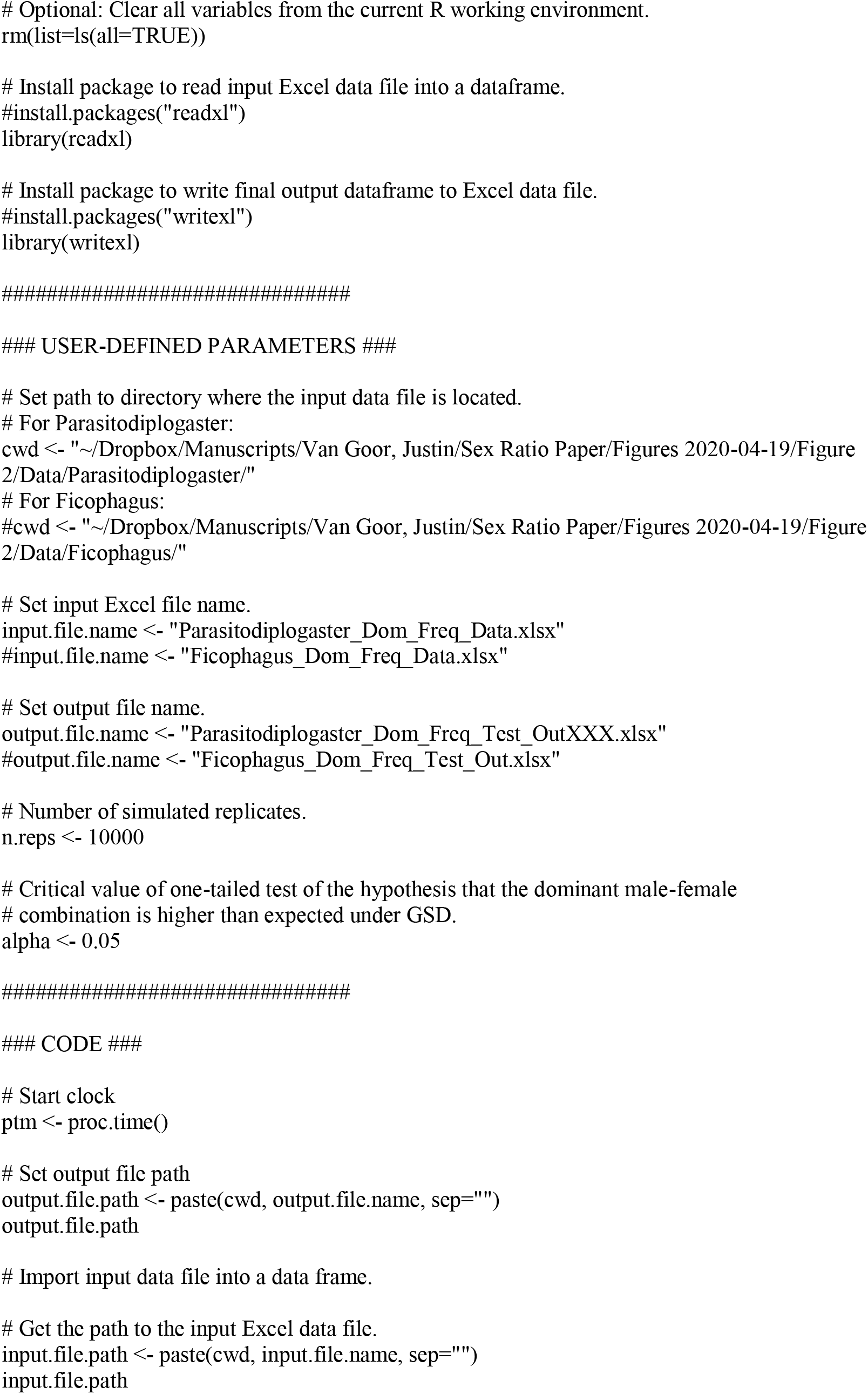
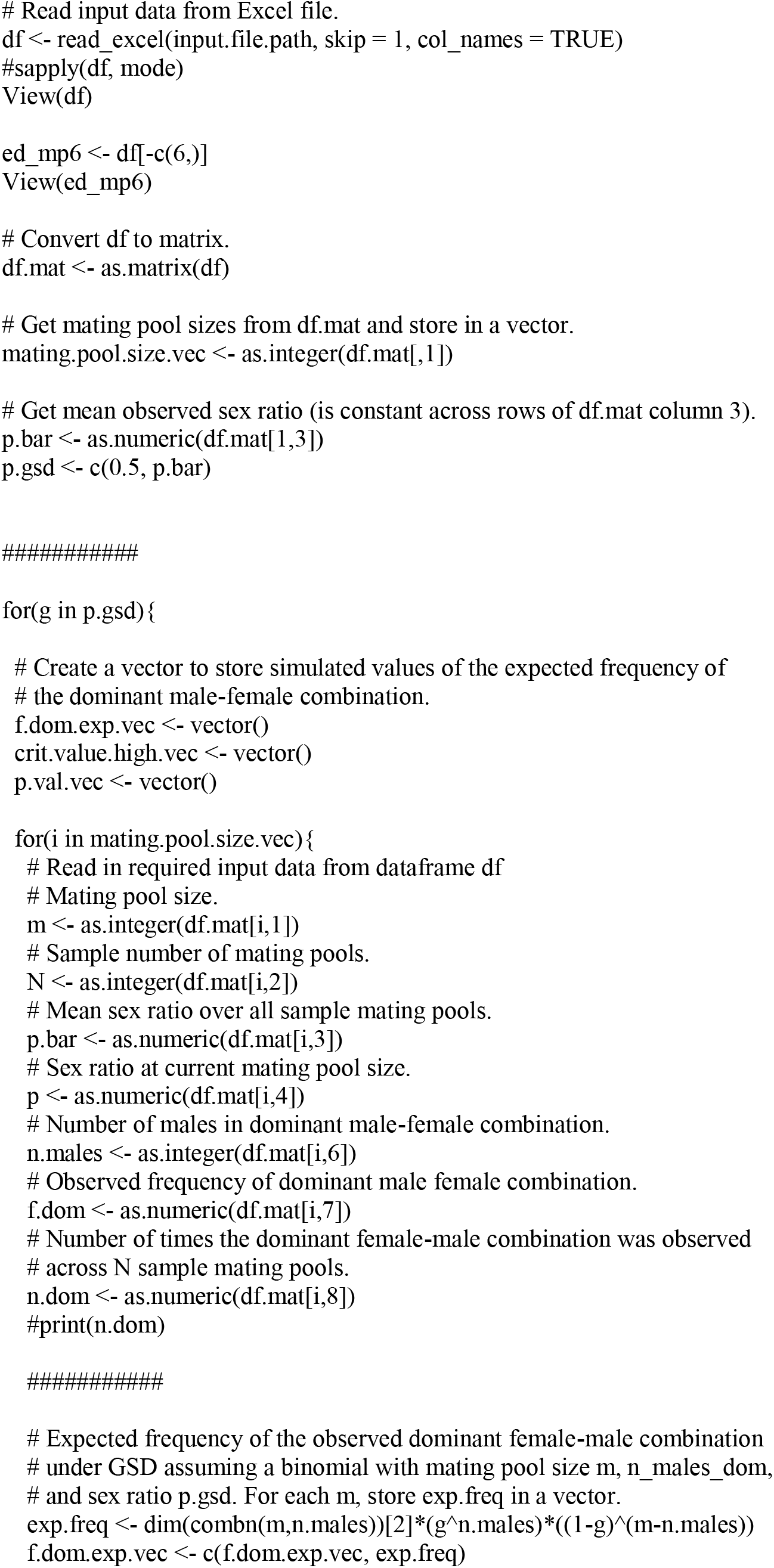
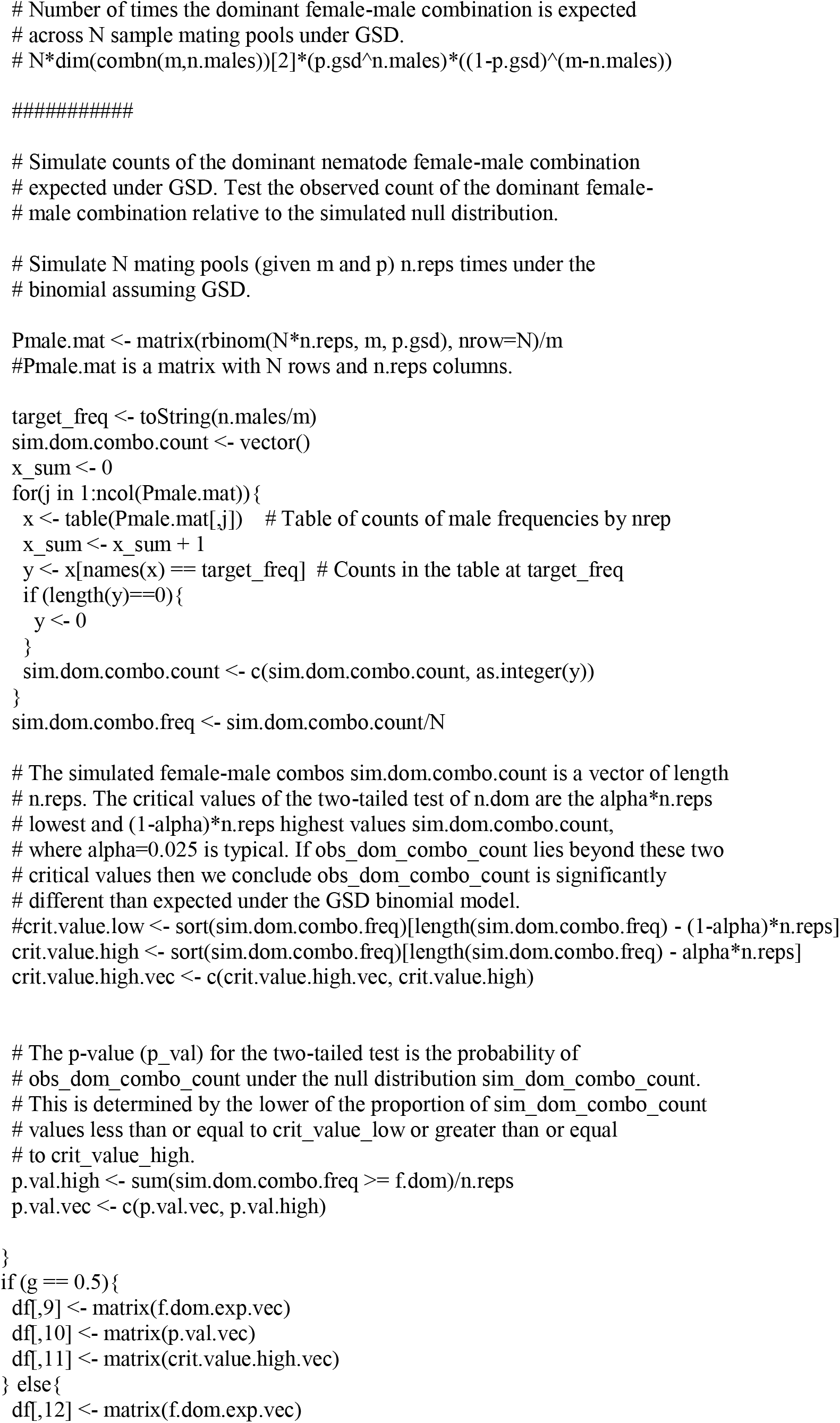
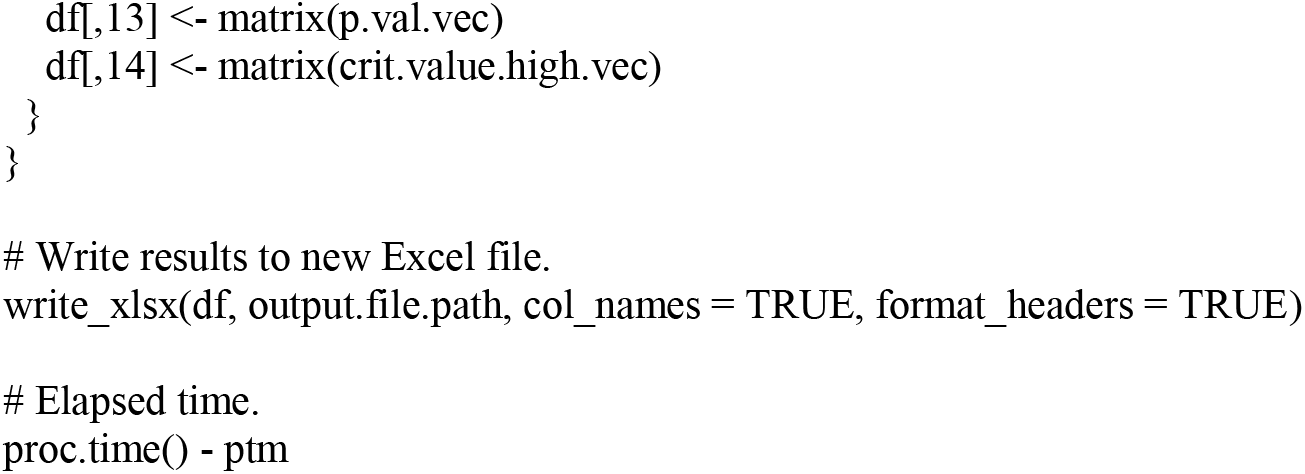

